# 8 Å structure of the nuclear ring of the *Xenopus laevis* nuclear pore complex solved by cryo-EM and AI

**DOI:** 10.1101/2021.11.10.468008

**Authors:** He Ren, Linhua Tai, Yun Zhu, Xiaojun Huang, Fei Sun, Chuanmao Zhang

## Abstract

The nuclear pore complex (NPC), one of the largest protein complexes in eukaryotes, serves as a physical gate to regulate nucleocytoplasmic transport. Here, we determined the 8 Å resolution cryo-electron microscopic (cryo-EM) structure of the nuclear ring (NR) from the *Xenopus laevis* NPC, with local resolutions reaching 4.9 Å. With the aid of AlphaFold2, we managed to build a pseudoatomic model of the NR, including the Y complexes and flanking components. In this most comprehensive and accurate model to date, the almost complete Y complex structure exhibits much tighter interaction in the hub region. Each NR asymmetric subunit contains two copies of Y complexes, one copy of Nup205 that connects the Y complexes to the neighbouring complex, one copy of ELYS that stabilizes the long arm region of the inner Y complex, and one copy of newly identified Nup93 that forms a bridge across the stems of Y complexes. These in-depth structural features represent a great advance in understanding the assembly of NPCs.

## INTRODUCTION

In eukaryotes, a double-layered membrane encloses a large organelle called the nucleus to separate genetic materials from the cytoplasm ^1^. The nuclear pore complex (NPC) fuses the inner and outer nucleus membrane (INM and ONM) to form the sole gateway mediating cargo transfer between the nucleoplasm and cytoplasm ^2,3^. NPCs are formed by approximately 30 different kinds of nucleoporins (Nups) with multiple copies in an eightfold symmetrical assembly. Each NPC contains approximately 550 proteins in fungi or approximately 1000 proteins in vertebrates ^4-6^. NPCs have a cylindrical appearance throughout their overall organization, where a central channel mediating bidirectional nucleocytoplasmic cargo transfer exists ^7,8^. Three scaffold rings, including an inner ring (IR) sandwiched by a nuclear ring (NR) and a cytoplasmic ring (CR) anchored onto the nuclear envelope (NE), provide bases for other functional parts, such as the cytoplasmic filament, nuclear basket, and permeability barrier ^9-12^.

As a major member of NPC scaffold rings, the NR is essential for building and maintaining NPC structures. It provides docking sites for the transportation of factors and cargos through the nuclear channel similar to pre-ribosomes and messenger ribonucleoproteins (mRNPS) ^13^ and stabilizes the nuclear basket substructure on the nucleoplasm side of the NPC. To date, several studies using cryo-electron tomography (cryo-ET) along with sub-tomo averaging (STA) have obtained the overall structures of the NR in the NPC ^9-12,14-19^, such as the study of Alexander et al., who reported the structure in 2015 ^15^ (PDB 5A9Q, aliased as 2015-model), and that of Lin et al., who reported the structure in 2016 ^20^ (aliased as 2016-model). The Y complex, also known as the Nup107-Nup160 complex, is the major building block of both the CR and NR and contains Nup85, Nup43, Nup37, Nup160, Nup96, Nup107, Nup133, Seh1 and Sec13 ^12,21-23^. These eight asymmetric units are arranged in a head-to-tail fashion to form the backbone of the NR, with Nup160 and Nup133 beta-propeller domains anchored onto the nuclear membrane ^12,15^. There is a question mark-shaped density sandwiched by the arms of two Y complexes, which has been identified as Nup188 or Nup205 ^15,20^. ELYS (<u>e</u>mbryonic <u>l</u>arge molecule derived from the <u>y</u>olk <u>s</u>ac, also known as Mel-28 or AHCTF1), an essential Nup for postmitotic NPC assembly, interacts with chromatin as well as some members of the Y complex ^15,24,25^. However, due to the lack of a high-resolution NR structure, protein components are recognized only at the domain level and rigidly docked by the X-ray structure.

In this study, we used cryo-electron microscopy (cryo-EM) single particle analysis (SPA) to collect data not only for NPCs on a flattened NE with stage tilting but also for the side-view NPCs on the edge of folded NE. Then, we determined the significantly improved cryo-EM map of the NR of the *Xenopus laevis* (*X. laevis*) NPC with an isotropic resolution of 8 Å and a local resolution reaching 4.9 Å. Meanwhile, with the aid of the recently emerged highly accurate protein structure prediction tool AlphaFold2 ^26^, we built the most complete pseudoatomic model of the NR, which includes Y complexes and multiple flanking components. With this significantly improved model, we identified novel structural features of Nups in the NR, including tight interactions in the Y complex hub, fruitful interactions between Nup205 and other Nups, detailed interactions between ELYS and Nup160, and the existence of Nup93 as a bridge across the stems of Y complexes. Our results provide an in-depth advance in our understanding of the structural features and assembly of NPCs.

## RESULTS

### Structure determination of the NR subunit of the *X. laevis* NPC

Considering that the NPCs on a flattened NE showed severe preferred orientation problems yielding anisotropic resolution of the cryo-EM map, we used a sample collection strategy according to previous structural studies for *X. laevis* NPCs^14^. Briefly, in addition to the data collection with the stage tilting angle set from 30 to 60 degrees, NPCs on the edge of a folded NE, which are shown in the side-view orientation, were especially selected and imaged (Extended Data Fig. 1). Application of this approach significantly improved Fourier space sampling and resulted in cryo-EM reconstructions with isotropic resolution (Extended Data Fig. 2). Three local masks were applied to cover the whole NR asymmetric unit to improve the local maps. The final average resolutions were 8.1 Å for the NR subunit region, 7.8 Å for the NR core region and 8.6 Å for the NR Nup133 region (Extended Data Fig. 1, 2), where the highest local resolution reached was 4.9 Å for the most rigid part of the NR subunit. According to the three-dimensional Fourier shell correlation (3D-FSC) estimates ^27^, all three reconstructions have sphericity scores better than 0.9 (0.91 to 0.94), suggesting no significant anisotropy in the reconstructions (Extended Data Fig. 2).

By using AlphaFold2 ^26^, we predicted highly accurate protein structures of all NPC Nups from *X. laevis* or *Xenopus tropicalis* (*X. tropicalis*). Then, with the help of molecular dynamics flexible fitting (MDFF) refinement and manual adjustment, we successfully modelled 21 components into a density map of the NR subunit (Supplementary Video 1), with reliable secondary structure details (Fig. 1A-B, Extended Data Fig. 3). It is worth noting that the refined model of each NR component against its corresponding local map is highly consistent with the model predicted by AlphaFOLD2, and only slight bending or domain shifts were found in some models (Extended Data Fig. 4), indicating that the structural predictions were highly accurate ^26^. The overall model-to-map resolution reached 9 Å, similar to the resolution estimated from two half maps (Extended Data Fig. 2D), confirming that no significant resolution anisotropy exists in our reconstructions ^27^. In our model of the NR subunit, a total of 15664 residues were built. Compared to a previously reported model for a human NPC, we extended the number of residues by approximately 82.6% compared to the NR in the 2015-model (8578 residues) and by approximately 41.6% compared to the NR in the 2016-model (11064 residues) ^15,20^. Since the structural features of β-propeller domains in Nups have been well established, these extensions are mainly located in α-helical regions ^1,2^.

**Figure 1.**
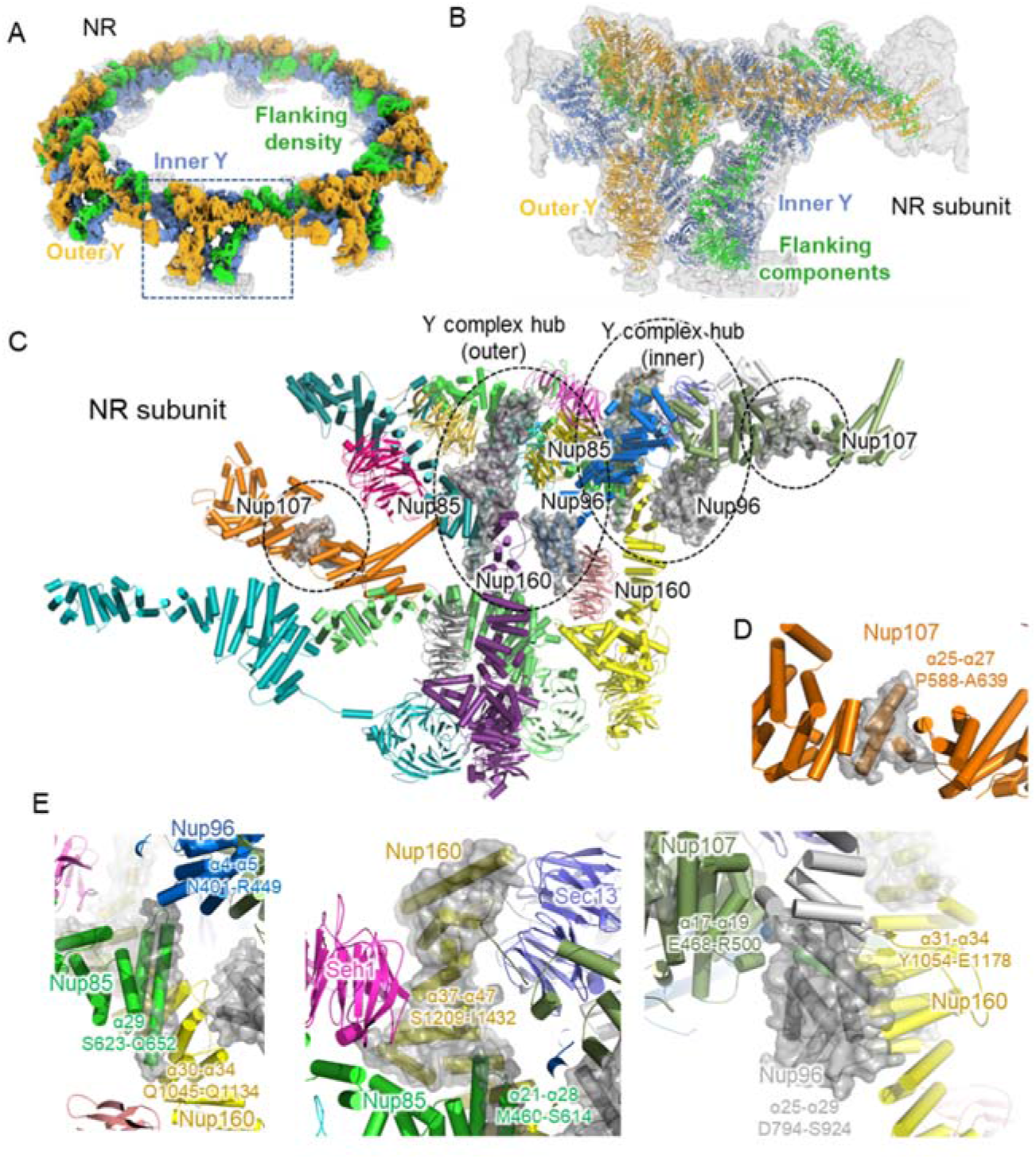
The more complete pseudoatomic model of the NR from the *X. laevis* oocyte NPC. **(A)** Overall view of the *X. laevis* NPC NR structure, displaying the inner and outer Y complexes in each asymmetric unit and densities other than the Y complexes. The inner Y complexes are presented in cornflower blue, the outer Y complexes are in orange, and the extra densities are in lime. **(B)** Model-map overlay of the NPC NR asymmetric unit. The map density is displayed in grey with transparency. Models of the inner Y complex, outer Y complex and extra densities are in the same colours as in **(A). (C)** Model display of two Y complexes in the NR asymmetric unit. All Nups are coloured differently. The major differences compared with the previous model are highlighted in the transparent grey surface. **(D)** Zoomed-in view of Nup107, displayed the same as in (C). (**E**) Zoomed-in view of the Y complex hub region, displayed the same as in (C). Important regions are labelled with the sequence and secondary structure numbers.

### The C-terminal domains (CTDs) of Nup85, Nup160 and Nup96 stabilize the Y complex hub

The outer diameter of the NR from the *X. laevis* NPC is 125 nm, while the inner diameter is 75 nm, both of which are consistent with previous reports ^12,14^. Two Y complexes lay in parallel, with a slight shift relative to one another, to form the backbone of the asymmetric unit of the NR (Fig. 1A). Most Nups of the Y complex have been well studied in both fungi and vertebrates. Nup85, Seh1 and Nup43 form the short arm of the Y complex, Nup160 and Nup37 form the long arm, and Sec13, Nup96, Nup107 and Nup133 form the stem, where the stem base covers Sec13, Nup96 and the N-terminal domain (NTD) of Nup107 and the stem tip covers the C-terminus of Nup107 and Nup133 (Fig. 1B-C) ^12,15^. Two domain invasion motifs (DIMs) exist in Nup85-Seh1 and Nup96-Sec13 to form complete β-propeller structures ^21,28,29^. These typical structures and features are all confirmed in our map and models (Extended Data Fig. 3).

In our most complete NR model from the *X. laevis* NPC, we identified several new features in the Y complex. In addition to the completion of the previously broken Nup107 structure, we clarified the important interaction hub connecting the two arms and stem base involving the CTDs of Nup85, Nup160 and Nup96 (Fig. 1C and D). For Nup85, we modelled a total of 29 α-helices, and S623-Q652 form the last long α-helix, which makes contact with Nup96 (α4-α5, N401-R449) and Nup160 (α30-α34, Q1045-Q1134) (Fig. 1E) ^20^. For Nup160, compared with 36 previously identified α-helices in the 2016-model, we extended it to 47α-helices until reaching the C-terminus. The α37-α47 helices at the Nup160 CTD (S1209-I1432) form the core structure of the Y complex hub by making contact with Seh1, Nup85 (α21-α28, M460-S614), and Sec13 (Fig. 1E). For Nup96, which starts with the DIM in Sec13, we modelled all 29 α-helices in total, with the last 5 α-helices (D794-S924) making contact with Nup107 (α17-α19, E468-R500) and Nup160 (α31-α34, Y1054-E1178) (Fig. 1E). Overall, Nup85, Nup160 and Nup96 come together at their CTDs, forming a hub structure to maintain the overall Y-shaped structure.

### One copy of Nup205 resides in the *X. laevis* NR subunit

In the cryo-ET map of a human NPC (EMD-3103), a question mark-shaped density was found to be sandwiched between two Y complexes in both the CR and NR ^15^. This density can be fitted well with the crystal structures of both Nup188 and Nup192 (homologues of Nup205) from *Chaetomium thermophilum* (ctNup188 and ctNup192) ^20,30-33^ because Nup188 and Nup205 share highly similar domain distributions (Fig. 2A). Some studies proposed that this density was more likely to be Nup188, since cross-linking mass spectrometry (XL-MS) revealed cross-linked pairs between Nup85 and Nup188 ^12,15^. Other studies suggested that ctNup192 has a long helix (named the tower helix) in its middle domain (MID), which can fit the density well, but the corresponding region in Nup188 lacks structural information ^20^. Most recently, the high-resolution structure of the *X. laevis* NPC attributed the question mark-shaped densities in the CR to Nup205, mainly according to the density of the tower helix ^34^.

**Figure 2.**
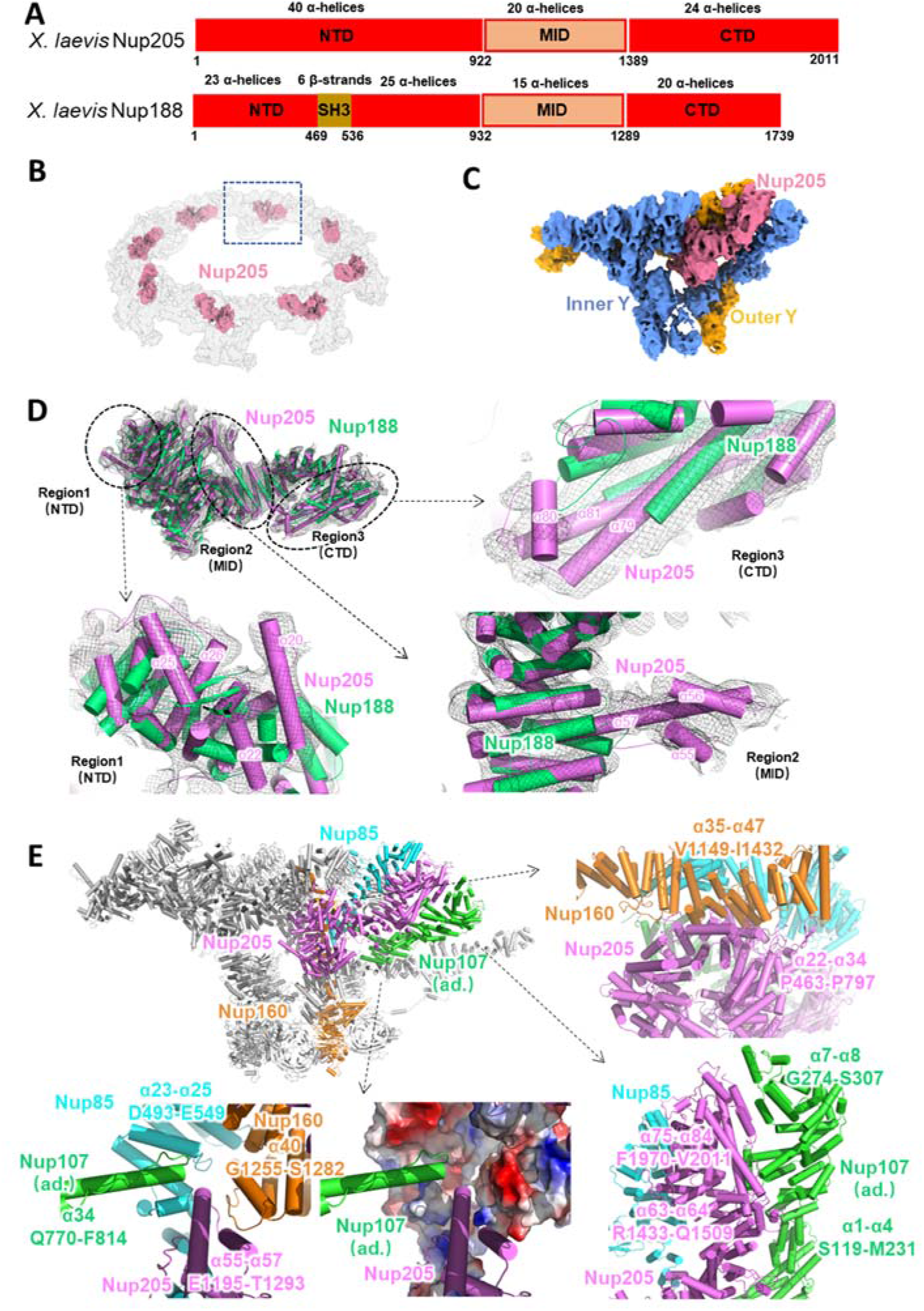
Nup205, instead of Nup188, connects the Y complexes within the same NR subunit and between adjacent subunits. **(A)** Domain assignment of *X. laevis* Nup205 and Nup188. **(B)** Location of Nup205 in the NR. **(C)** Location of Nup205 in the NR asymmetric unit. **(D)** Model comparison between *X. laevis* Nup205 and Nup188; the cryo-EM density is shown as a black mesh. Major differences are enlarged and displayed in detail. **(E)** Interaction of Nup205 with the outer Nup85 and outer Nup160 in the same NR subunit and the inner Nup107 in an adjacent (ad.) subunit. All discussed Nups are coloured differently. Interaction sites are labelled in detail.

To identify this component more accurately, we subtracted the densities of both Y complexes from our NR subunit, cropped the only remaining single copy of the question mark-shaped density (Fig. 2B and C), and built the model of Nup205 or Nup188 based on the structure predicted by AlphaFOLD2. In our models, there are 78 helices for Nup188 and 84 helixes for Nup205 (Fig. 2D). Overall, the Nup205 structure can fill this cropped density, and most of its helices fit the map very well, but Nup188 leaves densities unfilled in at least 3 local regions, which is easy to understand since Nup205 is 272aa longer than Nup188 in *X. laevis*. In the first region at the NTD of Nup205, four α helices (α20, α22, α25 and α26) match well with the local density. However, in this region, Nup188 has an SH3 domain forming several β-strands and could hardly match the map ^32^ (Fig. 2D). In the second region at the MID of Nup205, the most distinguishable structure of the tower helix (α57), wrapped by two shorter α-helices (α55 and α56), fits well with the map density, but Nup188 has only several short helices in this region that are not long enough to occupy it (Fig. 2D). In the third region at the CTD of Nup205, two long helices (α79 and α81) with a short helix (α80) that connects them can fill the local map density, but Nup188’s short helices fail to do so (Fig. 2D). Therefore, it can be concluded that there should be only one copy of Nup205 in the *X. laevis* NR subunit.

The complete Nup205 model in the NR subunit shows rich interactions with nearby NR components (Fig. 2E). In the NTD, the 22^nd^ to 34^th^ α-helices of Nup205 (P463-P497) connect to the CTD of outer Nup160 (α35 to α47, V1149 to I1432) at the Y complex hub (Fig. 2E). In the MID, the tower helix of Nup205 (α55-α57, E1195-T1293), together with the finger helix of Nup107 (α34, Q770-F814) in the adjacent NR subunit, is inserted into the grooves formed by Nup85 (α23-α25, D493-E549) and Nup160 (α40, G1255-S1282). These four proteins are grouped together in this local region to form a strong interaction network (Fig. 2E). In the CTD, the 63^rd^ and 64^th^ α-helices and 75^th^ to 84^th^ α-helices of Nup205 form tight contacts with the 1^st^ to 4^th^ α-helices and 7^th^ and 8^th^ α-helices of inner Nup107 in the adjacent NR subunit, respectively (Fig. 2E). These results suggest that Nup205 plays an important role in stabilizing the head-to-tail connection of asymmetric units of the NR ^20^.

### The ELYS NTD interacts with four adjacent Nups in the *X. laevis* NR

ELYS in the *X. laevis* NR contains an N-terminal β-propeller domain and a subsequent α-solenoid domain in the NTD and an unstructured domain in the CTD (Fig. 3A). A previous study found that ELYS anchors to the NE through its β-propeller domain and interacts with Nup160 through its α-helical domain; thus, ELYS was thought to be one of the Y complex Nups that localized in the NR ^15,24,25^. Stoichiometry research and the cryo-EM structure of the human NPC NR suggested that in each NR subunit, there are two copies of ELYS binding to both the inner and outer Nup160 ^6,15^. However, in this study, according to the high-resolution and isotropic cryo-EM map of the *X. laevis* NR, we could find only one copy of the ELYS density situated on the convex side of the inner Nup160 in each NR asymmetric unit (Fig. 3B and C). Even when using a mask large enough to cover the possible existing ELYS density on the outer Nup160 during data processing, we still failed to identify the second copy of ELYS in the NR subunit. Therefore, it is possible that ELYS has different copy numbers between human and *X. laevis*.

**Figure 3.**
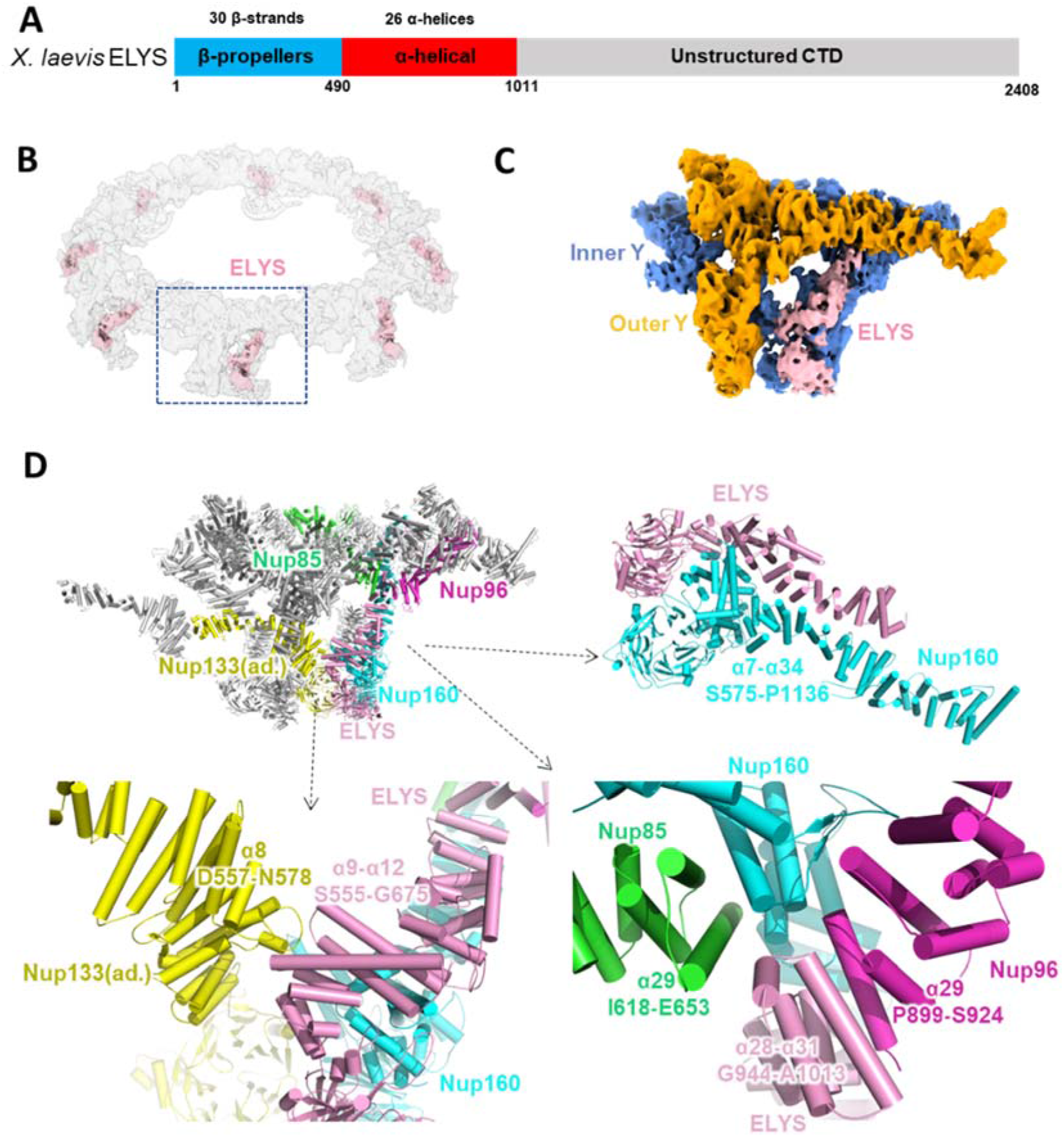
ELYS connects Nup160, Nup96 and Nup85 within the same NR subunit and Nup133 in an adjacent subunit. **(A)** Domain assignment of *X. laevis* ELYS. **(B)** Location of ELYS in the NR. **(C)** Location of ELYS in the NR asymmetric unit. **(D)** Interaction of ELYS with the convex inner Nup160 and inner Y complex hub in the same NR subunit and the inner Nup133 in the adjacent subunit. All discussed Nups are coloured differently. Interaction sites are labelled in detail.

In our NR structure, all the domains in the ELYS NTD are modelled well in terms of the map density (Extended Data Fig. 3B). It is obvious that ELYS has extensive and tight interactions in both the β-propeller domain and α-solenoid domain with the inner Nup160 (β-propeller domain and α7-α34) (Fig. 3D), suggesting that during the ELYS-mediated NPC assembly process, Nup160 could be easily recruited by ELYS. In addition to the inner Nup160, ELYS also interacts with the inner Nup85, inner Nup96 and inner Nup133 (Fig. 3D). The 9^th^ to 12^th^ α-helices (S555-G675) of ELYS come close to the 8^th^ α-helix (D557-N578) of the inner Nup133 in the adjacent NR subunit, stabilizing the head-to-tail connection of the NR subunits. The C-terminus of the ELYS NTD, α28-α31 helices (G944-A1013), participated in the formation of the inner Y complex hub. In addition to the inner Nup160, this region interacts with the 29^th^ α-helix of the inner Nup85 and the 29^th^ helix of the inner Nup96 (Fig. 3D). It is worth noting that due to the interaction between ELYS and Nup96, the density of the inner Nup96’s C-terminus seems more rigid than that of the outer Nup96’s C-terminus while playing with the map threshold level. This also suggests that ELYS is missing in the outer Y complex in the NR of *X. laevis* NPC.

Therefore, our model exhibits the 3D conformations of ELYS in the NR asymmetric unit and shows its interactions with four surrounding NR components, i.e., Nup160, Nup133, Nup85 and Nup96. These results show that ELYS significantly contributes to the structural stability of the NR and provide clues for understanding the NPC postmitotic assembly process introduced by ELYS.

### Nup93 bridges the adjacent Y complex in the *X. laevis* NR subunit

Nup93 (Nic96 in fungi) (Fig. 4A) has been identified as a building block of the IR complex (IRC) through interactions with channel nucleoporin heterotrimers (CNTs) and Nup205/Nup188 ^20,30^. Usually, Nup93 is not believed to be localized in the NR and involved in NR assembly. However, after assignment of other densities in the NR subunit, we noticed that there is a distinct bridge-like density on the stem region of the two Y complexes (Fig. 4B and C). The unknown density at this location has also been depicted in the human NPC ^15^, but there is still no evidence of any Nups being assigned for this density. In our high-resolution NR map, this bridge-like density is divided into two distinct α-helical regions, whose centres are approximately 8 nm apart. The smaller density connects the inner Nup160 and outer Nup107, while the larger density forms an apparent α-solenoid domain to link the inner Nup96 and outer Nup107.

**Figure 4.**
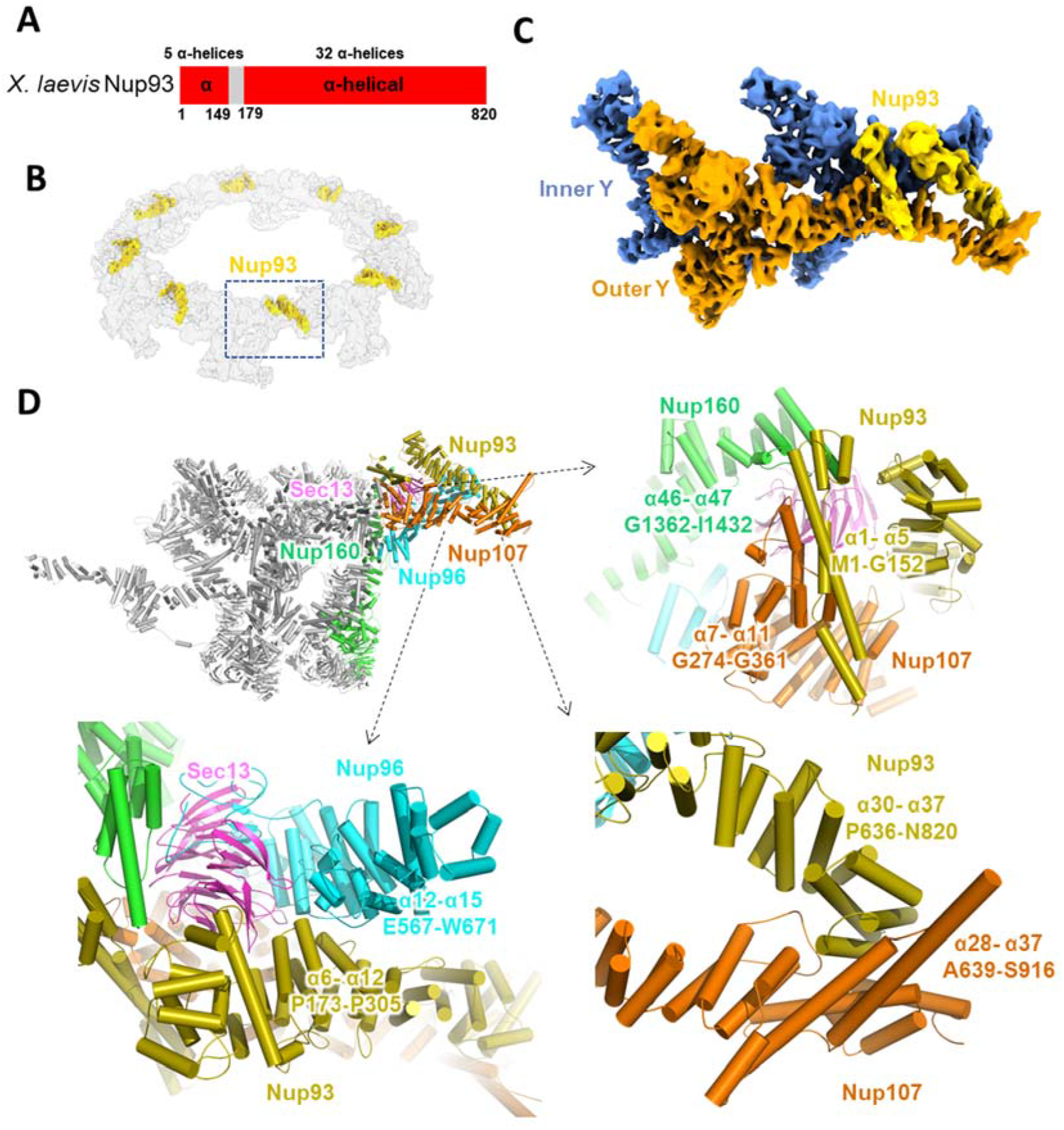
Nup93 acts as a bridge to connect the stems of the inner and outer Y complexes. **(A)** Domain assignment of *X. laevis* Nup93. **(B)** Location of Nup93 in the NR. **(C)** Location of Nup93 in the NR asymmetric unit. **(D)** Interaction of Nup93 with the inner Nup96, inner Nup160 and outer Nup107. All discussed Nups are coloured differently. Interaction sites are labelled in detail.

After trying to fit the predicted structures of all known Nups from *X. laevis*, we found that only Nup93 matched both densities well (Extended Data Fig. 3G, Supplementary Video 2). The model of Nup93 predicted by AlphaFOLD2 has two domains, a 5 α-helix NTD and a 32 α-helix CTD, which are connected by a long flexible loop. The special long 5^th^ helix in the Nup93 NTD, which has approximately 50 residues and is even longer than Nup205’s tower helix and Nup107’s finger helix ^35^, matches well with the smaller bridge-like density. The cross-correlation (CC) value of the model to map within the mask for Nup93 was calculated to be 0.71 by PHENIX ^36^.

The NTD of Nup93 (α1-α5, M1-G152) interacts with the CTD of the inner Nup160 (α46-α47, G1362-I1432) and outer Nup107 (α7-α11, G274-G361) to connect the stems of the two Y complexes (Fig. 4C). The connecting loop of Nup93 (aa 153-179) allows the separation of the NTD and CTD with a long distance of 8 nm. The Nup93 CTD forms a typical α-solenoid domain, whose 6^th^ to 12^th^ α-helices are in the proximity of the 12^th^ to 15^th^ α-helices of the inner Nup96, and the 30^th^ to 37^th^ α-helices connect to the 28^th^ to 37^th^ α-helices of the outer Nup107. Therefore, both the NTD and CTD of Nup93 play an essential role in connecting the most flexible parts of the two Y complexes in the stem region, thus maintaining the stability of the NR subunit.

## Discussion

Modelling the macromolecular complex according to a cryo-EM map with middle resolution has long been hindered by the lack of complete and reliable structural information for all the components. These components are sometimes too flexible or unstable in monomer form, which makes it very difficult to purify them and determine their high-resolution structures ^15,20^. Recently, the protein structure prediction method of AlphaFOLD2 has been proved to predict 3D structures with an accuracy level that no other programs have reached ^26^, which provides an important tool for modelling the complete structure of macromolecular complexes, such as the NPC. In this study, we found that all the structures of the *X. laevis* NR Nups predicted by AlphaFOLD2 matched the actual density map very well, and only a few additional refinements (MDFF, PHENIX.real_space_refine, manual refine, etc.) were needed to obtain a complete structural model. Some domain rotations or shifts were found after the refinement in some Nups, such as Nup96, Nup133, ELYS and Nup93 (Extended Data Fig. 4), which may be due to conformational changes in monomer proteins during complex assembly. We believe that the structure prediction tools will be able to calculate the complex structure soon, which will greatly facilitate the structure determination of macromolecular complexes.

To correctly interpret the map density, the resolution anisotropy must be considered for samples with severe orientational problems, such as the NPC on a flattened NE ^14,34^. We have made great efforts in solving the NPC structure combined with 0-, 30-, 45-, and 60-degree tilted images, but the 3D-FSC estimates always suggested that the resolutions along the Z axis were only approximately 20 Å, while the resolutions along the other two axes could reach 6∼8 Å. This kind of anisotropic map made assignment of the secondary structure features difficult. Thus, in order to eliminate resolution anisotropy as far as possible, we introduced side-view NPC particles into the dataset and omitted 0-degree tilting images from the very beginning of image process, this strategy helped to solve this problem by compensating for imperfect Fourier space sampling, improving the Z axis resolution to better than 10 Å (Extended Data Fig. 2).

In contrast to vertebrate NPCs, NPCs in fungi exhibit less conservation in the overall shape of their outer rings. In each asymmetric unit of the NPC from *Chlamydomonas reinharadtii* (*C. reinharadtii*) or *Schizosaccharomyces pombe* (*S. pombe*), the CR has only one Y complex, and the NR has two Y complexes in each asymmetric unit, while in *Saccharomyces cerevisiae* (*S. cerevisiae*), both outer rings have only one Y complex in each asymmetric unit ^15,17,18^. Nup358 is located in the stem region of the CR, similar to Nup93 in the NR, and knockdown of Nup358 could cause a lack of the formation of two Y complexes in the NPC CR ^15,17^. This proved that Nup358 is necessary to maintain the two Y complex conformations in the CR. However, the role of Nup93 in the NR might be different from that of Nup358 in the CR. Among the NR maps in different species (Extended Data Fig. 5), we found that the Nup93 corresponding density was present only in higher organisms, such as vertebrates, and was missing in *S. pombe* and *C. reinharadtii*, the NPCs of which also had two Y complexes in each NR asymmetric unit ^15,17^. Therefore, Nup93 does not seem necessary to maintain the two Y complex conformations in the NR.

In all species of humans, toads, algae, and yeast (*S. pombe*), ELYS is present in the genome as a full-length protein, or in truncated form, or as a homologue protein ELY5 ^15,17,24,25^. ELYS has been proved to participate in Y complex hub formation and play a role in connecting adjacent Y complexes. In yeast, ELY5 has only α-helical regions and thus makes no contribution to anchoring the Y complex onto the NE but may play a similar role in stabilizing the Y complex hub. According to sequence alignment, ELY5 contains α-helical regions corresponding to the 13^th^ to 30^th^ α-helices of ELYS, which cover the major interaction sites of ELYS α-solenoid domain with respect to Nup160. ELYS is highly evolutionarily conserved in both structures and functions, indicating that the existence of ELYS in NPCs is essential for NPC assembly and the evolution of the NR from single-cell organisms to vertebrates.

In summary, we developed an improved approach to solve the cryo-EM map of the *X. laevis* NPC NR in an isotropic resolution of 8 Å, built an almost complete pseudoatomic model with the aid of AlphaFold2 to obtain an accurate secondary structure, and discovered novel structural details, providing a critical update to our understanding of the assembly of NPCs.

## Methods

### Sample preparation

African clawed toad *X. laevis* maintenance, oocyte isolation, and NE preparation for cryo-EM were carried out as described previously ^8,14,37^. Briefly, ovaries were removed from narcotized mature female *X. laevis* (Nasco, USA) with a brief wash in freshly prepared amphibian Ringer’s solution (111 mM NaCl, 1.9 mM KCl, 1.1 mM CaCl_2_, 2.4 mM NaHCO_3_), and developmental stage VI oocytes were transferred to ice-cold HEPES buffer (83 mM KCl, 17 mM NaCl, 10 mM HEPES, pH 7.5) for nuclear isolation. For side-view NPCs, the oocyte nucleus was gently penetrated and let it falls onto the grid. For top-view NPCs, the oocyte nuclei were isolated in HEPES buffer, applied to glow-discharged holey carbon grids (R2/1, 200 mesh, Au, Quantifoil, Germany), and the NE was spread onto the grid by fine glass needles. After careful washing in HEPES buffer, the NE on the grid was cross-linked with 0.15% glutaraldehyde in HEPES buffer on ice for 10 min. Then the grid was blotted and vitrified by plunge freezing into liquid ethane by Vitrobot Mark IV (Thermo Fisher Scientific, USA) at 4 °C and 100% humidity.

The animal experiments were performed in the Laboratory Animal Center of Peking University in accordance with the National Institutes of Health Guide for the Care and Use of Laboratory Animals and according to guidelines approved by the Institutional Animal Care and Use Committee at Peking University.

### Cryo-EM data acquisition

During the application of the NE onto the grid, for flattened NE, the CR side was kept always facing the carbon film of the grid. Prior to data collection, we first separated the grids with flattened NE into two groups: one group with the carbon film facing onto the C-clip of FEI AutoGrid and another group with the carbon film facing in the opposite direction. All grids were screened using a Talos Arctica 200 kV cryo-electron microscope (Thermo Fisher Scientific, USA). Then, 8745 micrographs were collected using a Titan Krios G2 300 kV cryo-electron microscope (Thermo Fisher Scientific, USA) operated in EF-TEM mode with a nominal magnification of 64,000X. The calibrated physical pixel size on the specimen was 2.24 Å. For flattened NR specimens, the stage tilting angles were set to 0, 30, 45 and 60 degrees. For the 0/30-degree tilting angles, the total exposure dose was set to 60 e^-^/Å^2^, with an exposure time of 21.5 seconds and 0.5 seconds per frame. For the 45-degree tilting angle, the total exposure dose was set to 80 e^-^/Å^2^, with an exposure time of 41 or 28.5 seconds for two different sessions, 0.5 seconds per frame. For the 60-degree tilting angle, the total exposure dose was set to 100 e^-^/Å^2^, with an exposure time of 36 or 35 seconds for two different sessions, 0.5 seconds per frame.

For side-view particles, the total exposure dose was set to 100 or 120 e^-^/Å^2^ with an exposure time of 28.5 or 34.5 seconds. The movies were acquired by a Gatan K2 Summit direct detection camera equipped with a GIF Quantum energy filter (Gatan Company, USA) with a silt width of 20 eV operated in super-resolution mode. SerialEM with in-house scripts was used for data collection with the defocus value set between 1.0 to 4.0 μm ^38,39^.

### SPA image processing

The super-resolution movies were first subjected to motion correction using MotionCor2 with a binning level of 2 in Fourier space, and dose weighting was also performed during this process ^40^. Since tilted images required accurate defocus value estimation on a per particle basis, particle picking was performed prior to CTF estimation. Particles were auto-picked using RELION-3.0 with subsequent manual inspection ^41^. A total of 85,065 full NPC particles were selected. As during data collection of side-view NPCs the stage tilt was set to be 0-degree, and 0-degree tilt images of flattened NE sample were not introduced into image processing, we use 0-degree image to refer to side-view images from this point on. For 0-degree images, the defocus of each picked NPC was estimated by Gctf using its per particle defocus estimation function ^42^. For other tilting images, the defocus of each selected NPC was estimated by goCTF or Warp ^43,44^.

The selected NPC particles were first extracted with a box size of 216 pixels at a binning level of 4, which resulted in a pixel size of 8.96 Å (Extended Data Fig. 1). Since NPCs on the same flattened NE pointing to approximately same direction, we first ran global search for each particle correspond to each tilting angles, using the previously reported NPC map (EMDB entry EMD-3103) low-pass filtered to 60 Å as the initial reference, with 8-fold symmetry applied ^15^. Then we ran statistics of tilt angles for each dataset. For the dataset with carbon film pointing to C-clip of AutoGrid, their tilt angles were smaller than 90 degrees, and were approximately equal to stage tilting angles. For those datasets with carbon film pointing to C-ring, their tilt angles approximately equaled to 90 degrees add with stage tilt angles. With this prior information on the tilting angles and the relative orientation of the grid, we were able to assign a prior tilt angle of 30/45/60/120/135/150 degrees for each particle. For side-view particles, the initial tilt angle was set to be 90 degrees.

The refinements were performed using RELION-3.0 with the local search strategy applied. First, we ran 100 iterations of 3D classification with K = 1 using the previously reported NPC map (EMD-3103) low-pass filtered to 60 Å as the initial reference, and 8-fold symmetry was applied, the tilt angle search range was restricted to 3 degrees with respect to previous value for each iteration ^15^. Then, all particles were merged to run another round of 3D classification with K = 1 for 30 iterations in 8-fold symmetry, using the NPC map (EMD-3103) low-pass filtered to 60 Å as reference. Then, we docked the previously reported model of the NR (PDB entry code: 5A9Q) into the resulting map and segmented out the NR region from the whole NPC using Chimera ^45^. Based on the segmented map, we generated a local mask covering solely the NR region. Using this mask and the output star file from the previous round of 3D classification, we performed auto-refinement to obtain a 20 Å resolution map of the NR with the 8-fold symmetry applied.

With the refined shifts and orientations of NR of the NPC particles, we re-extracted particles with a box size of 400 pixels and binned pixel size of 4.48 Å. Using a similar strategy shown above, we reconstructed the cryo-EM map of the NR at a resolution of 18 Å. With this better resolution, we achieved a more accurate determination of each NR asymmetric unit. We used Chimera to measure the relative coordinates of one asymmetric unit of the NR and generated a symmetry expanded particle star file with updated defocus corresponding to each NR asymmetric unit by using a modified version of a block-based reconstruction script ^46^. Then, we re-extracted particles with a box size of 200 pixels at a binning level of 2, pixel size 4.48 Å. Another round of auto-refinement was performed using the NR asymmetric unit mask, yielding the cryo-EM map of the NR asymmetric unit at a resolution of around 10 Å.

With the refined shifts and orientations of each asymmetric unit, we re-extracted particles with a box size of 320 pixels, binning level of 1 and pixel size of 2.24 Å. We first performed a reconstruction to ensure that the predetermined parameters were correct and to generate a better mask containing one asymmetric unit of the NR. Then, another round of auto-refinement yielded a map at a resolution of 9 Å. Next, we ran Bayesian polishing for all particles, then separated the polished particles into different groups based on individual stage tilt angles. Next, we ran four different 3D classification jobs for 30-, 45-, 60-degree dataset, where 45-degree stage tilting dataset was further separated into 45-degree and 135-degree datasets with T values of 20, 20, 20, 8, respectively, 120-degree datasets was not subjected to 3D classification due to small number of particles. Particles corresponding to the best class were selected, and merged with side-view particles and 120 degree tilting particles, then an auto-refinement job of these particles was done using RELION 3.0. After this round of refinement, the output particles, map, and mask were transferred into CryoSPARC to do local refinement which reached a final resolution of 8.1 Å ^41,47^. A rather stable NR core region was identified by investigating the local resolution distribution of this 8.1 Å map using RELION-3.0, and the mask covering this region only was created ^41^. Then a local refinement of the NR core region was performed using the same data as 8.1 Å map, which resulted in a final resolution of 7.8 Å ^47^. A similar strategy was applied to the NR Nup133 region of the NR and resulted in a resolution of 8.6 Å. The validation of map and model quality of this research was done using 3D-FSC and Phenix ^27,48^ (Extended Data Fig. 2).

### Modeling of NPC NR

The full version of AlphaFold2 was downloaded from GitHub and installed as instructed with all database downloaded ^26^. All the structures of NPC Nups from *X. laevis* or *X. tropicalis* (Nup160 and Nup96), were predicted by AlphaFOLD2 using the recommended calculation parameters. The value of Max_template_hits was set to 20, Relax_energy_tolerance was set to 2.39, Relax_stiffness was set to 10, Relax_max_outer_iterations was set to 20. For each Nup, a total of 5 relaxed structures were predicted, and the prediction with the highest confidence was selected as the starting model for the next refinement.

Then we performed stepwise MDFF simulations to refine each Nup model according to the corresponding local cryo-EM density in NR subunit. A timestep of 1 fs was used throughout the simulation. Langevin dynamics were adopted at a temperature of 310 K. The equilibration step for energy minimization was performed on the initial model for 1000 steps before the refinement run. The refinement runs were performed for 3000 ps, which corresponds to 3,000,000 simulation steps, and the gridForceScale values were gradually increased from 0.3 to 0.7 during the refinement. All simulations were performed using CHARMM36m forcefields ^49^. Electrostatic calculations were treated with particle mesh Ewald (PME). A cutoff of 12 Å was chosen for short-range van der Waals interactions. NAMD ^50^ was used as the MD engine throughout all simulations.

All the NR components were assembled in COOT ^51^ for manually adjustment according to the overall density map of NR subunit. Then the whole model of NR subunit was refined using PHENIX.real_space_refine ^36^. Data collection statistics and refinement statistics are given in Extended Data Table 1. All figures in this study were generated by PyMol, Chimera and ChimeraX ^45,52^.

## Data Availability

The Electron Microscopy Database (EMD) accession codes of the NR subunit region, NR core region and NR Nup133 region are EMD-31891, EMD-31892, EMD-31893, respectively. The Protein Data Bank (PDB) accession code of the model of the NR asymmetric unit is 7VCI.

## ACKNOWLEDGEMENTS

We thank all other members of the Fei Sun and Chuanmao Zhang laboratories for their help. We would also like to thank the Center for Biological Imaging (CBI), Institute of Biophysics, Chinese Academy of Science for cryo-EM work, and Boling Zhu, Xujing Li Gang Ji, Jiashu Xu and Guoliang Yin for their help with cryo-EM data collection; the Facilities Cores at National Center for Protein Sciences and the cryo-EM and TEM platforms at the College of Life Sciences of Peking University for cryo-electron microscopy and TEM; and Ning Gao, Zhenxi Guo, Guopeng Wang, Yingchun Hu, Xia Pei and Bo Shao for their help with cryo-EM and TEM experiments.

This work was equally supported by grants from Ministry of Science and Technology of China (2017YFA0504700 to FS and 2016YFA0500201 to CMZ), the Strategic Priority Research Program of the Chinese Academy of Sciences (XDB 37040102 to FS), and National Natural Science Foundation of China (31830020 to FS, 31520103906 to CMZ). This work was also supported by grants from the National Science Fund for Distinguished Young Scholars (31925026 to FS), National Natural Science Foundation of China (31430051 to CMZ) and National Key Research and Development Program of China (2016YFA0100501 to CMZ and 2018YFA0901102 to YZ).

## AUTHOR CONTRIBUTIONS

F. S. and C. Z. conceived the project and designed the experiments. L. T., H. R. and X. H. performed cryo-EM experiments. L. T. and Y. Z. performed cryo-EM data processing. H. R. and L. T. participated in the preparation and screening of cryo-EM samples. Y. Z. performed the modeling and simulation-based refinement. L. T. and Y. Z. analyzed the data and wrote the manuscript, which was substantially revised by F. S. and C. Z.

## References

1 Lin, D. H. & Hoelz, A. The Structure of the Nuclear Pore Complex (An Update). Annu Rev Biochem 88, 725–783, doi:10.1146/annurev-biochem-062917-011901 (2019).

2 Hoelz, A., Debler, E. W. & Blobel, G. The structure of the nuclear pore complex. Annu Rev Biochem 80, 613–643, doi:10.1146/annurev-biochem-060109-151030 (2011).

3 Hampoelz, B., Andres-Pons, A., Kastritis, P. & Beck, M. Structure and Assembly of the Nuclear Pore Complex. Annu Rev Biophys 48, 515–536, doi:10.1146/annurev-biophys-052118-115308 (2019).

4 Rout, M. P. et al. The yeast nuclear pore complex: composition, architecture, and transport mechanism. J Cell Biol 148, 635–651, doi:10.1083/jcb.148.4.635 (2000).

5 Cronshaw, J. M., Krutchinsky, A. N., Zhang, W., Chait, B. T. & Matunis, M. J. Proteomic analysis of the mammalian nuclear pore complex. J Cell Biol 158, 915–927, doi:10.1083/jcb.200206106 (2002).

6 Ori, A. et al. Cell type-specific nuclear pores: a case in point for context-dependent stoichiometry of molecular machines. Mol Syst Biol 9, 648, doi:10.1038/msb.2013.4 (2013).

7 Hinshaw, J. E., Carragher, B. O. & Milligan, R. A. Architecture and design of the nuclear pore complex. Cell 69, 1133–1141, doi:10.1016/0092-8674(92)90635-p (1992).

8 Akey, C. W. & Radermacher, M. Architecture of the Xenopus nuclear pore complex revealed by three-dimensional cryo-electron microscopy. J Cell Biol 122, 1–19, doi:10.1083/jcb.122.1.1 (1993).

9 Beck, M. et al. Nuclear pore complex structure and dynamics revealed by cryoelectron tomography. Science 306, 1387–1390, doi:10.1126/science.1104808 (2004).

10 Beck, M., Lucic, V., Forster, F., Baumeister, W. & Medalia, O. Snapshots of nuclear pore complexes in action captured by cryo-electron tomography. Nature 449, 611–615, doi:10.1038/nature06170 (2007).

11 Maimon, T., Elad, N., Dahan, I. & Medalia, O. The human nuclear pore complex as revealed by cryo-electron tomography. Structure 20, 998–1006, doi:10.1016/j.str.2012.03.025 (2012).

12 Bui, K. H. et al. Integrated structural analysis of the human nuclear pore complex scaffold. Cell 155, 1233–1243, doi:10.1016/j.cell.2013.10.055 (2013).

13 Delavoie, F., Soldan, V., Rinaldi, D., Dauxois, J. Y. & Gleizes, P. E. The path of pre-ribosomes through the nuclear pore complex revealed by electron tomography. Nat Commun 10, 497, doi:10.1038/s41467-019-08342-7 (2019).

14 Eibauer, M. et al. Structure and gating of the nuclear pore complex. Nat Commun 6, 7532, doi:10.1038/ncomms8532 (2015).

15 von Appen, A. et al. In situ structural analysis of the human nuclear pore complex. Nature 526, 140–143, doi:10.1038/nature15381 (2015).

16 Kim, S. J. et al. Integrative structure and functional anatomy of a nuclear pore complex. Nature 555, 475–482, doi:10.1038/nature26003 (2018).

17 Mosalaganti, S. et al. In situ architecture of the algal nuclear pore complex. Nat Commun 9, 2361, doi:10.1038/s41467-018-04739-y (2018).

18 Allegretti, M. et al. In-cell architecture of the nuclear pore and snapshots of its turnover. Nature 586, 796–800, doi:10.1038/s41586-020-2670-5 (2020).

19 Zhang, Y. et al. Molecular architecture of the luminal ring of the Xenopus laevis nuclear pore complex. Cell Res 30, 532–540, doi:10.1038/s41422-020-0320-y (2020).

20 Lin, D. H. et al. Architecture of the symmetric core of the nuclear pore. Science 352, aaf1015, doi:10.1126/science.aaf1015 (2016).

21 Hsia, K. C., Stavropoulos, P., Blobel, G. & Hoelz, A. Architecture of a coat for the nuclear pore membrane. Cell 131, 1313–1326, doi:10.1016/j.cell.2007.11.038 (2007).

22 Kampmann, M. & Blobel, G. Three-dimensional structure and flexibility of a membrane-coating module of the nuclear pore complex. Nat Struct Mol Biol 16, 782–788, doi:10.1038/nsmb.1618 (2009).

23 Kelley, K., Knockenhauer, K. E., Kabachinski, G. & Schwartz, T. U. Atomic structure of the Y complex of the nuclear pore. Nat Struct Mol Biol 22, 425–431, doi:10.1038/nsmb.2998 (2015).

24 Bilokapic, S. & Schwartz, T. U. Structural and functional studies of the 252 kDa nucleoporin ELYS reveal distinct roles for its three tethered domains. Structure 21, 572–580, doi:10.1016/j.str.2013.02.006 (2013).

25 Bilokapic, S. & Schwartz, T. U. Molecular basis for Nup37 and ELY5/ELYS recruitment to the nuclear pore complex. Proc Natl Acad Sci U S A 109, 15241–15246, doi:10.1073/pnas.1205151109 (2012).

26 Jumper, J. et al. Highly accurate protein structure prediction with AlphaFold. Nature 596, 583–589, doi:10.1038/s41586-021-03819-2 (2021).

27 Tan, Y. Z. et al. Addressing preferred specimen orientation in single-particle cryo-EM through tilting. Nat Methods 14, 793–796, doi:10.1038/nmeth.4347 (2017).

28 Brohawn, S. G., Leksa, N. C., Spear, E. D., Rajashankar, K. R. & Schwartz, T. U. Structural evidence for common ancestry of the nuclear pore complex and vesicle coats. Science 322, 1369–1373, doi:10.1126/science.1165886 (2008).

29 Debler, E. W. et al. A fence-like coat for the nuclear pore membrane. Mol Cell 32, 815–826, doi:10.1016/j.molcel.2008.12.001 (2008).

30 Stuwe, T. et al. Architecture of the fungal nuclear pore inner ring complex. Science 350, 56–64, doi:10.1126/science.aac9176 (2015).

31 Stuwe, T., Lin, D. H., Collins, L. N., Hurt, E. & Hoelz, A. Evidence for an evolutionary relationship between the large adaptor nucleoporin Nup192 and karyopherins. Proc Natl Acad Sci U S A 111, 2530–2535, doi:10.1073/pnas.1311081111 (2014).

32 Andersen, K. R. et al. Scaffold nucleoporins Nup188 and Nup192 share structural and functional properties with nuclear transport receptors. Elife 2, e00745, doi:10.7554/eLife.00745 (2013).

33 Sampathkumar, P. et al. Structure, dynamics, evolution, and function of a major scaffold component in the nuclear pore complex. Structure 21, 560–571, doi:10.1016/j.str.2013.02.005 (2013).

34 Huang, G. et al. Structure of the cytoplasmic ring of the Xenopus laevis nuclear pore complex by cryo-electron microscopy single particle analysis. Cell Res 30, 520–531, doi:10.1038/s41422-020-0319-4 (2020).

35 Boehmer, T., Jeudy, S., Berke, I. C. & Schwartz, T. U. Structural and functional studies of Nup107/Nup133 interaction and its implications for the architecture of the nuclear pore complex. Mol Cell 30, 721–731, doi:10.1016/j.molcel.2008.04.022 (2008).

36 Liebschner, D. et al. Macromolecular structure determination using X-rays, neutrons and electrons: recent developments in Phenix. Acta Crystallogr D Struct Biol 75, 861–877, doi:10.1107/S2059798319011471 (2019).

37 Allen, T. D. et al. A protocol for isolating Xenopus oocyte nuclear envelope for visualization and characterization by scanning electron microscopy (SEM) or transmission electron microscopy (TEM). Nat Protoc 2, 1166–1172, doi:10.1038/nprot.2007.137 (2007).

38 Mastronarde, D. N. Automated electron microscope tomography using robust prediction of specimen movements. J Struct Biol 152, 36–51, doi:10.1016/j.jsb.2005.07.007 (2005).

39 Wu, C., Huang, X., Cheng, J., Zhu, D. & Zhang, X. High-quality, high-throughput cryo-electron microscopy data collection via beam tilt and astigmatism-free beam-image shift. J Struct Biol 208, 107396, doi:10.1016/j.jsb.2019.09.013 (2019).

40 Zheng, S. Q. et al. MotionCor2: anisotropic correction of beam-induced motion for improved cryo-electron microscopy. Nat Methods 14, 331–332, doi:10.1038/nmeth.4193 (2017).

41 Zivanov, J. et al. New tools for automated high-resolution cryo-EM structure determination in RELION-3. Elife 7, doi:10.7554/eLife.42166 (2018).

42 Zhang, K. Gctf: Real-time CTF determination and correction. J Struct Biol 193, 1–12, doi:10.1016/j.jsb.2015.11.003 (2016).

43 Su, M. goCTF: Geometrically optimized CTF determination for single-particle cryo-EM. J Struct Biol 205, 22–29, doi:10.1016/j.jsb.2018.11.012 (2019).

44 Tegunov, D. & Cramer, P. Real-time cryo-electron microscopy data preprocessing with Warp. Nat Methods 16, 1146–1152, doi:10.1038/s41592-019-0580-y (2019).

45 Pettersen, E. F. et al. UCSF Chimera--a visualization system for exploratory research and analysis. J Comput Chem 25, 1605–1612, doi:10.1002/jcc.20084 (2004).

46 Zhu, D. et al. Pushing the resolution limit by correcting the Ewald sphere effect in single-particle Cryo-EM reconstructions. Nat Commun 9, 1552, doi:10.1038/s41467-018-04051-9 (2018).

47 Punjani, A., Rubinstein, J. L., Fleet, D. J. & Brubaker, M. A. cryoSPARC: algorithms for rapid unsupervised cryo-EM structure determination. Nat Methods 14, 290–296, doi:10.1038/nmeth.4169 (2017).

48 Afonine, P. V. et al. Towards automated crystallographic structure refinement with phenix.refine. Acta Crystallogr D Biol Crystallogr 68, 352–367, doi:10.1107/S0907444912001308 (2012).

49 Huang, J. et al. CHARMM36m: an improved force field for folded and intrinsically disordered proteins. Nat Methods 14, 71–73, doi:10.1038/nmeth.4067 (2017).

50 Phillips, J. C. et al. Scalable molecular dynamics with NAMD. J Comput Chem 26, 1781–1802, doi:10.1002/jcc.20289 (2005).

51 Emsley, P., Lohkamp, B., Scott, W. G. & Cowtan, K. Features and development of Coot. Acta Crystallogr D Biol Crystallogr 66, 486–501, doi:10.1107/S0907444910007493 (2010).

52 Goddard, T. D. et al. UCSF ChimeraX: Meeting modern challenges in visualization and analysis. Protein Sci 27, 14–25, doi:10.1002/pro.3235 (2018).

