## Supplemental Data for "8 Å structure of the nuclear ring of the *Xenopus laevis* nuclear pore complex solved by cryo-EM and AI"

### Extended data figures and tables

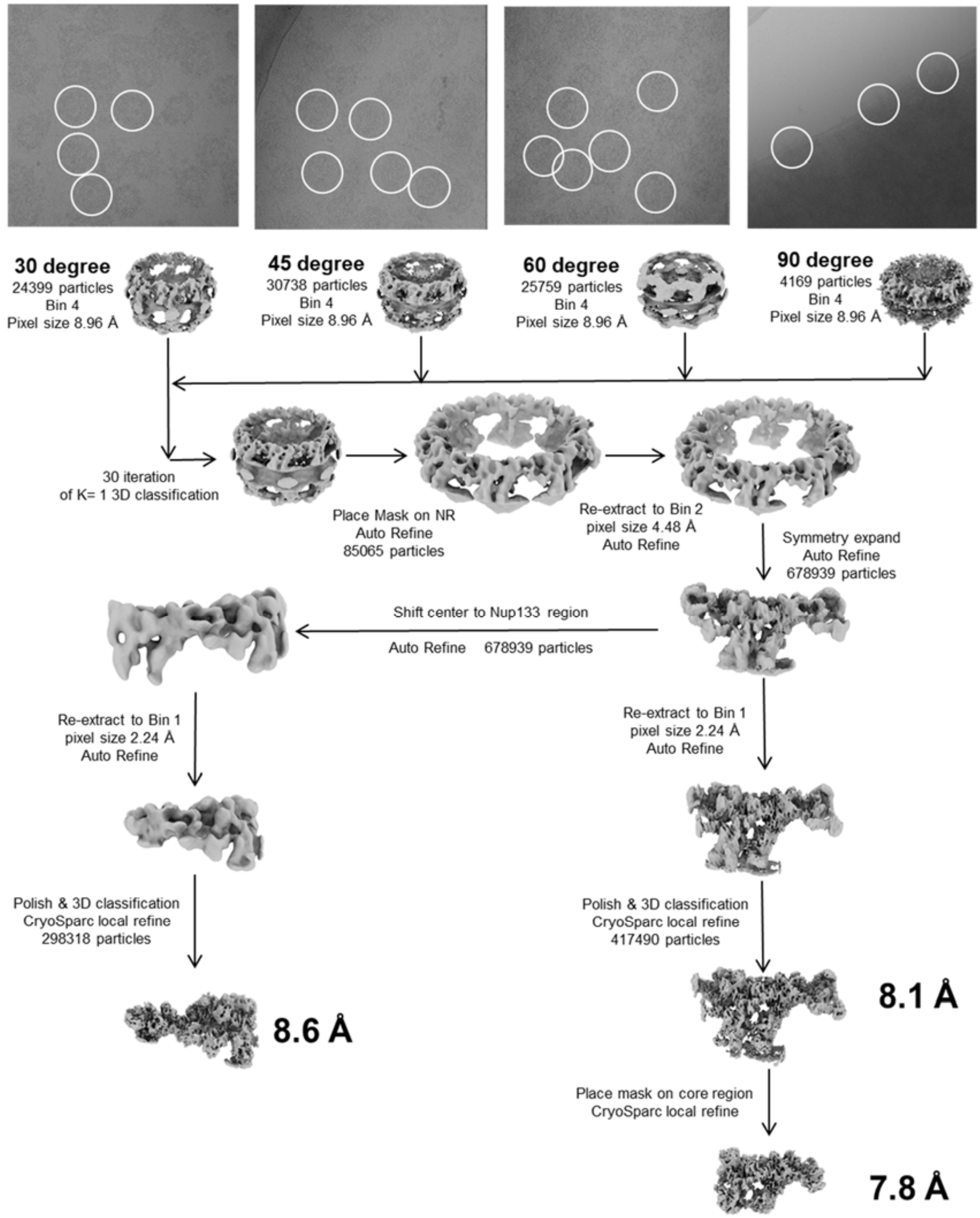


**Extended Data Figure 1. Data processing workflow.**

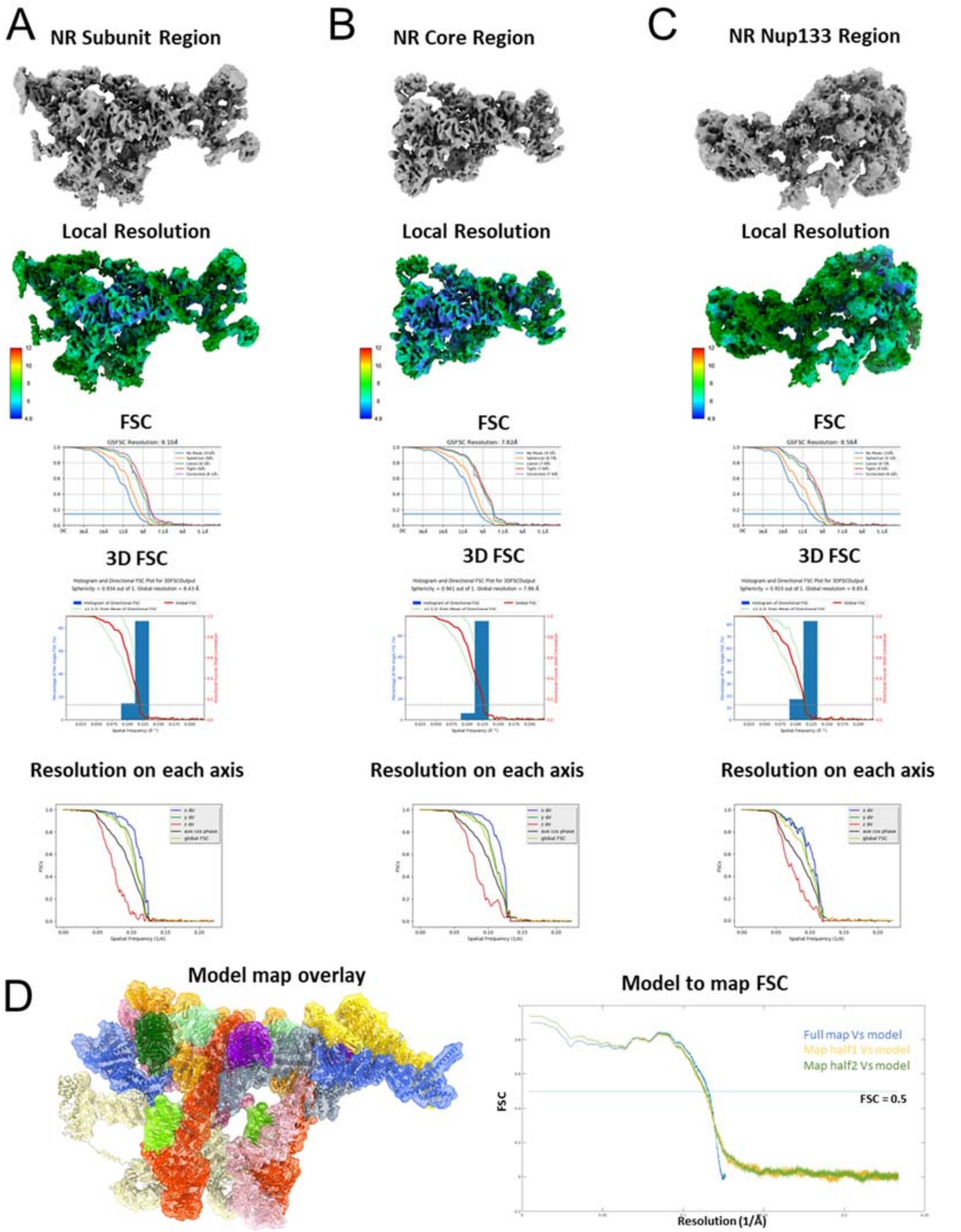


**Extended Data Figure 2. Assessments of cryo-EM maps. (A, B, C)** Map display, local resolution estimation, FSC, 3D-FSC and directional FSC of the NR subunit region, NR core region and NR Nup133 region. **(D)** Map model overlay of the NR asymmetric unit, and map versus model FSC estimation.

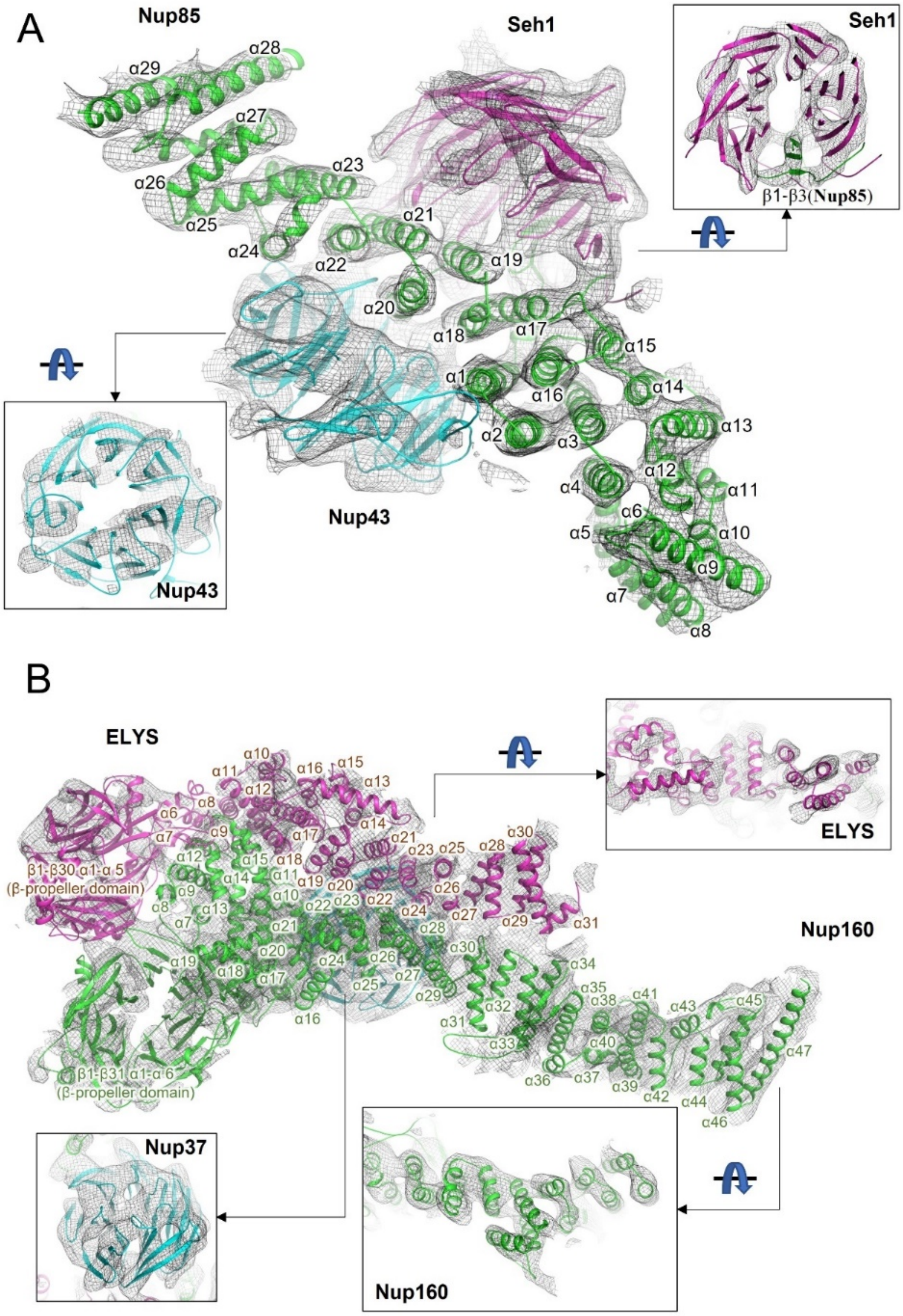

A
Nup85
Seh1
Seh1
β1-β3(Nup85)
Nup43
Nup43
B
ELYS
ELYS
β1-β30 α1-α 5
(β-propeller domain)
Nup160
β1-β31 α1-α 6
(β-propeller domain)
Nup37
Nup160

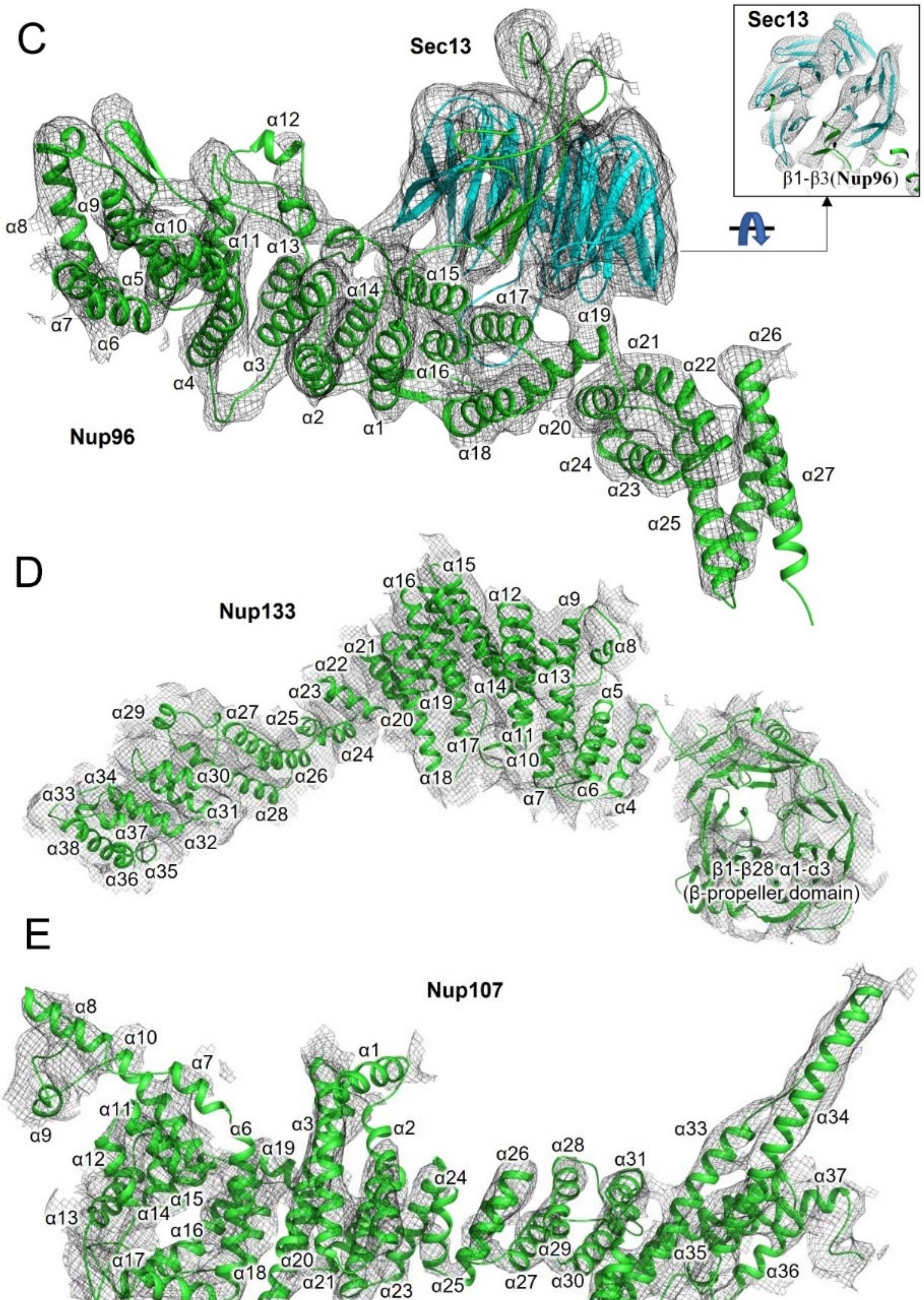
C
Sec13
Nup96
Sec13
β1-β3(Nup96)
D
Nup133
β1-β28 α1-α3
(β-propeller domain)
E
Nup107

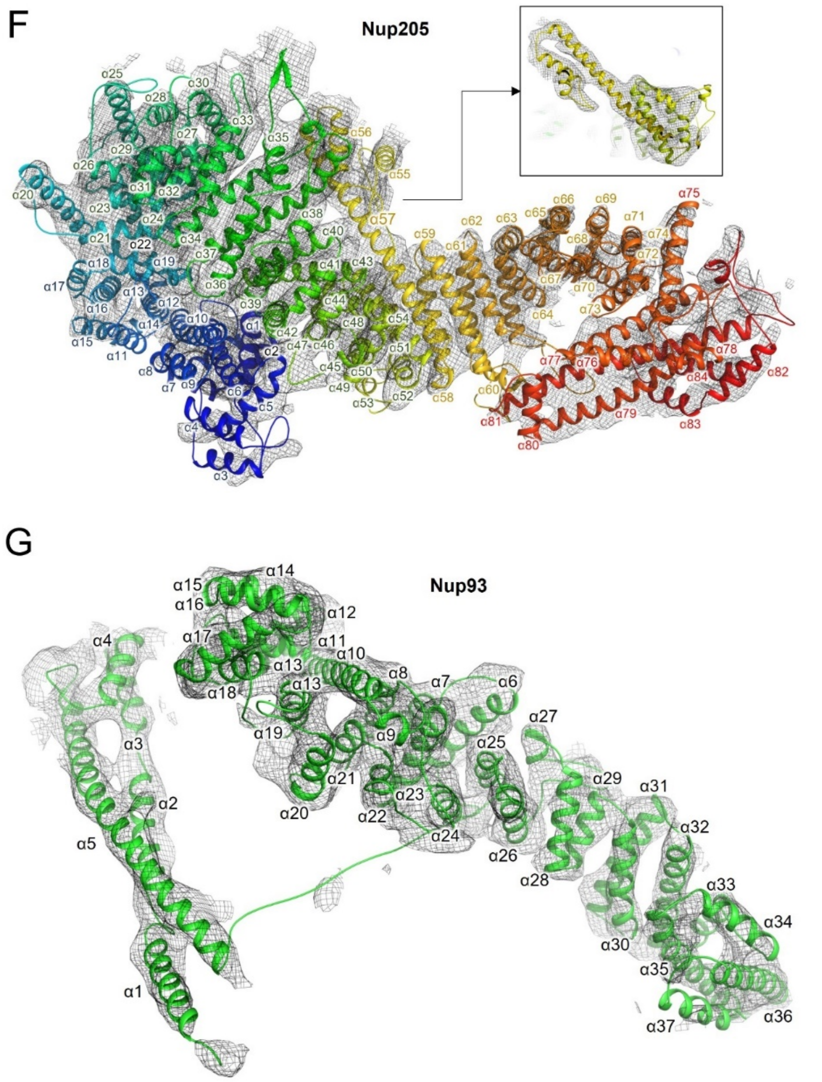


**Extended Data Figure 3. Model building quality and secondary structure assignment of the NR asymmetric unit. (A-G)** Model building quality of Nup85, Seh1, Nup43, ELYS, Nup160, Sec13, Nup96, Nup133, Nup107, Nup205 and Nup93.

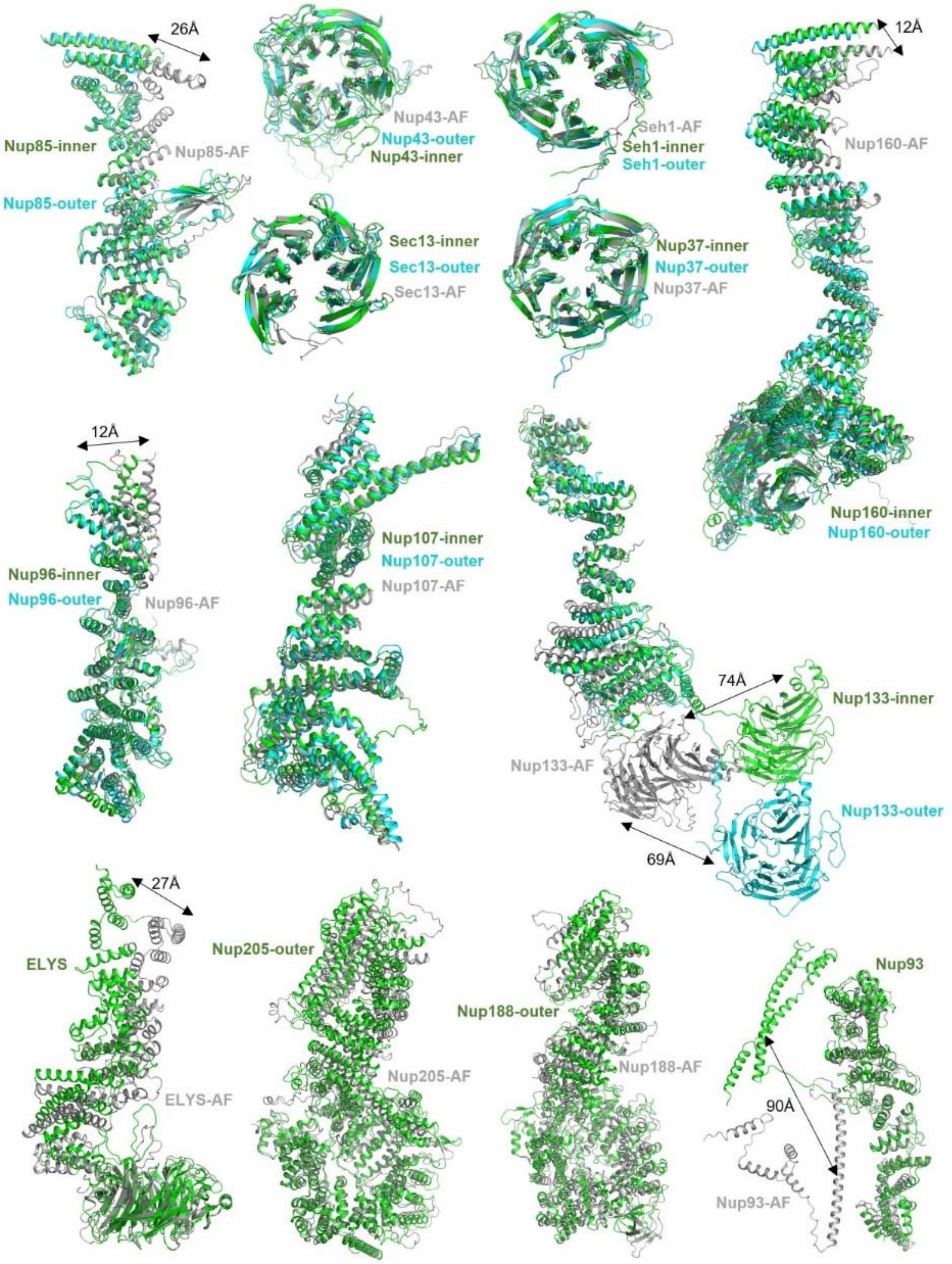


**Extended Data Figure 4. Comparisons of the refined structure and predicted structure by AlphaFold2 (AF) of NR Nups from the *X. laevis* NPC.**

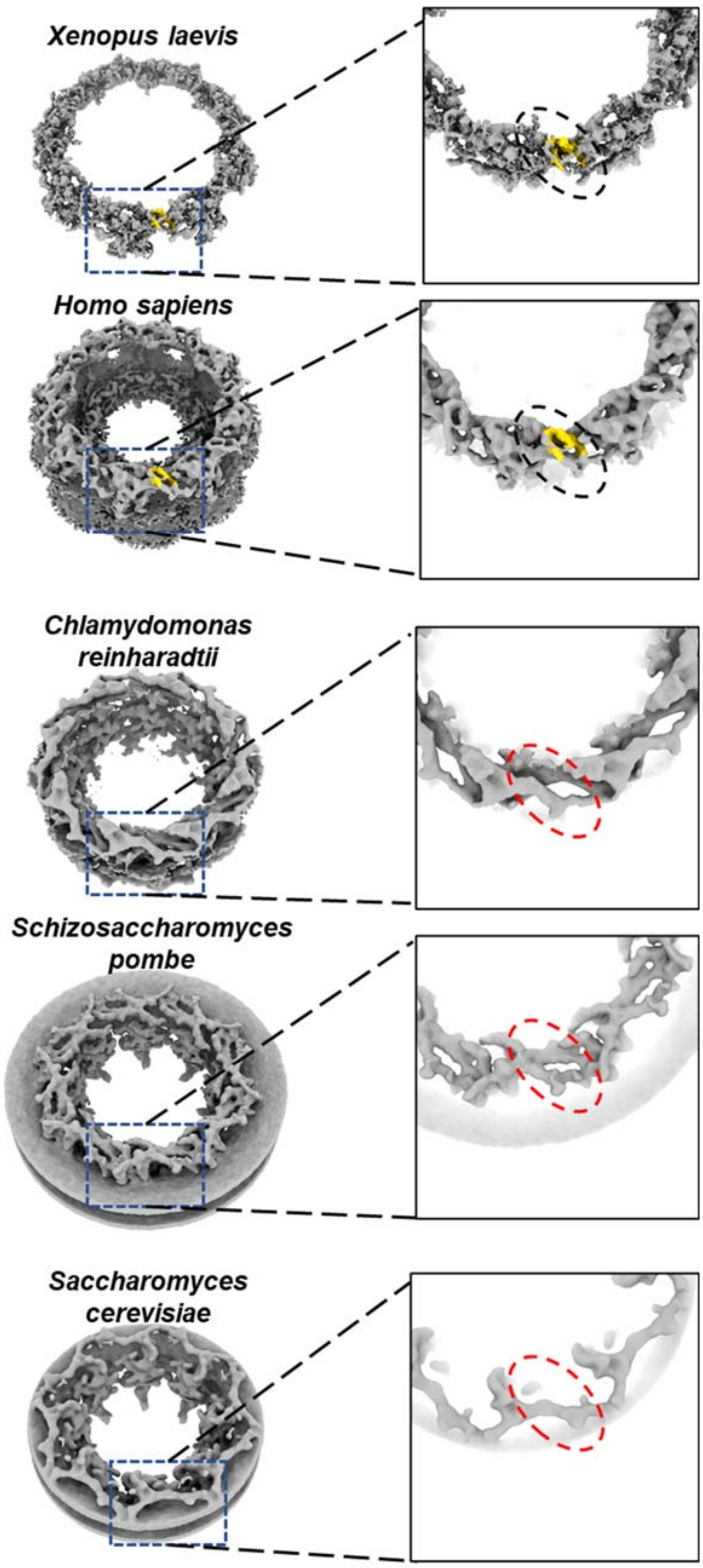


**Extended Data Figure 5. Comparisons of bridge-like densities in the NPC NR for different species.**

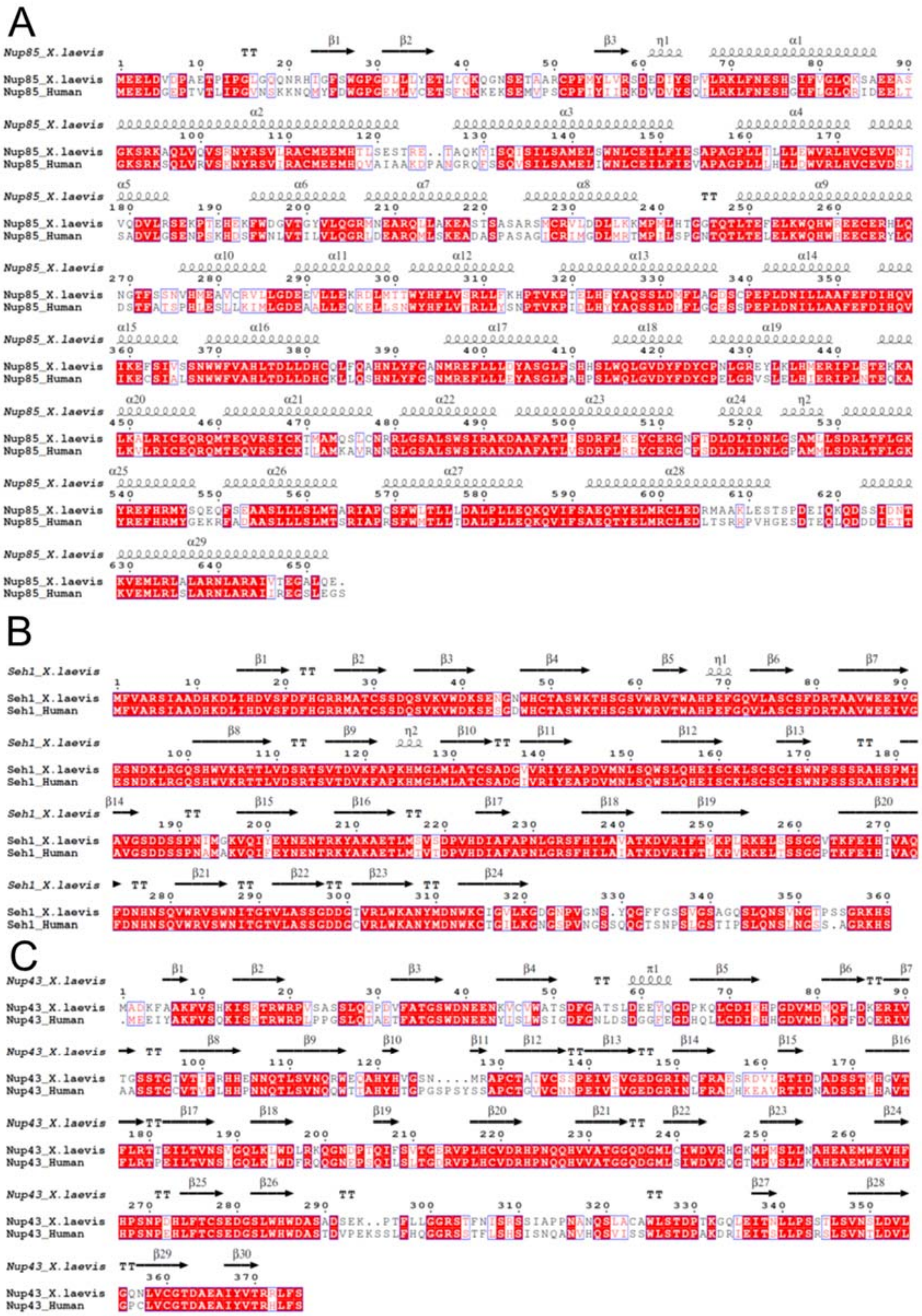
A
Nup85_X.laevis
Nup85_Human
B
Seh1_X.laevis
Seh1_Human
C
Nup43_X.laevis
Nup43_Human

D

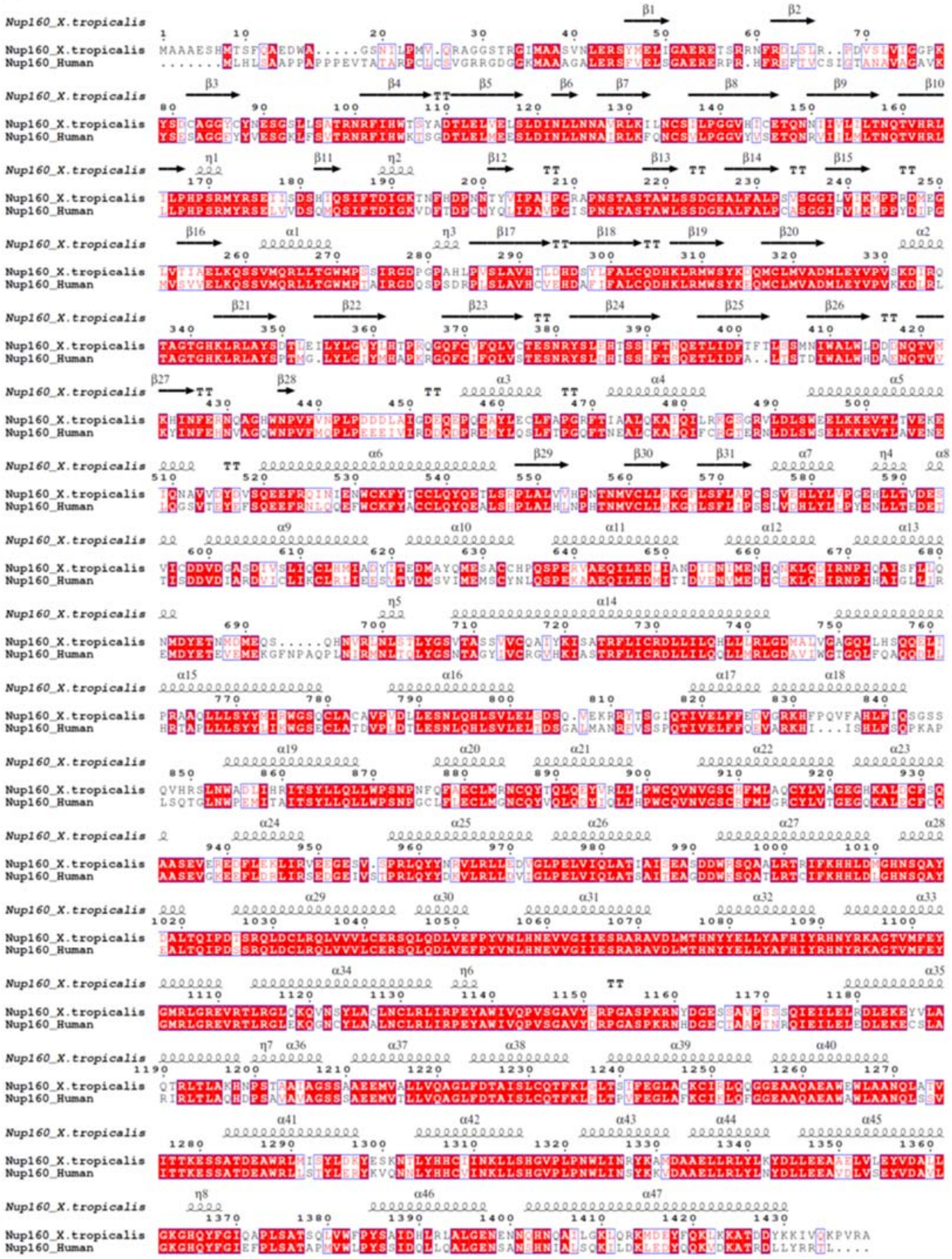

E

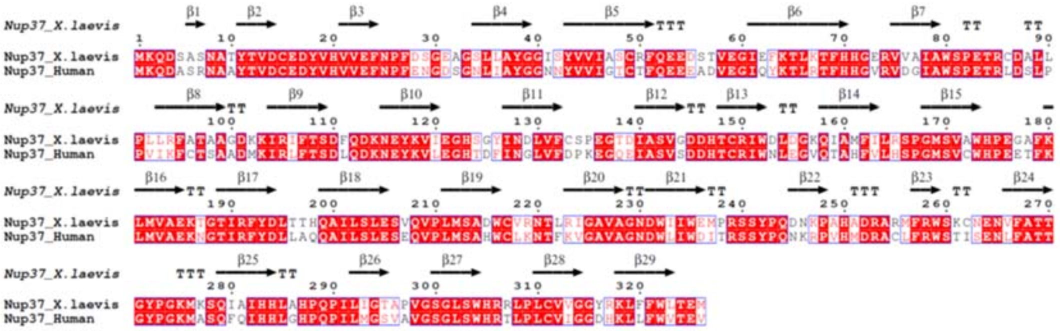


F

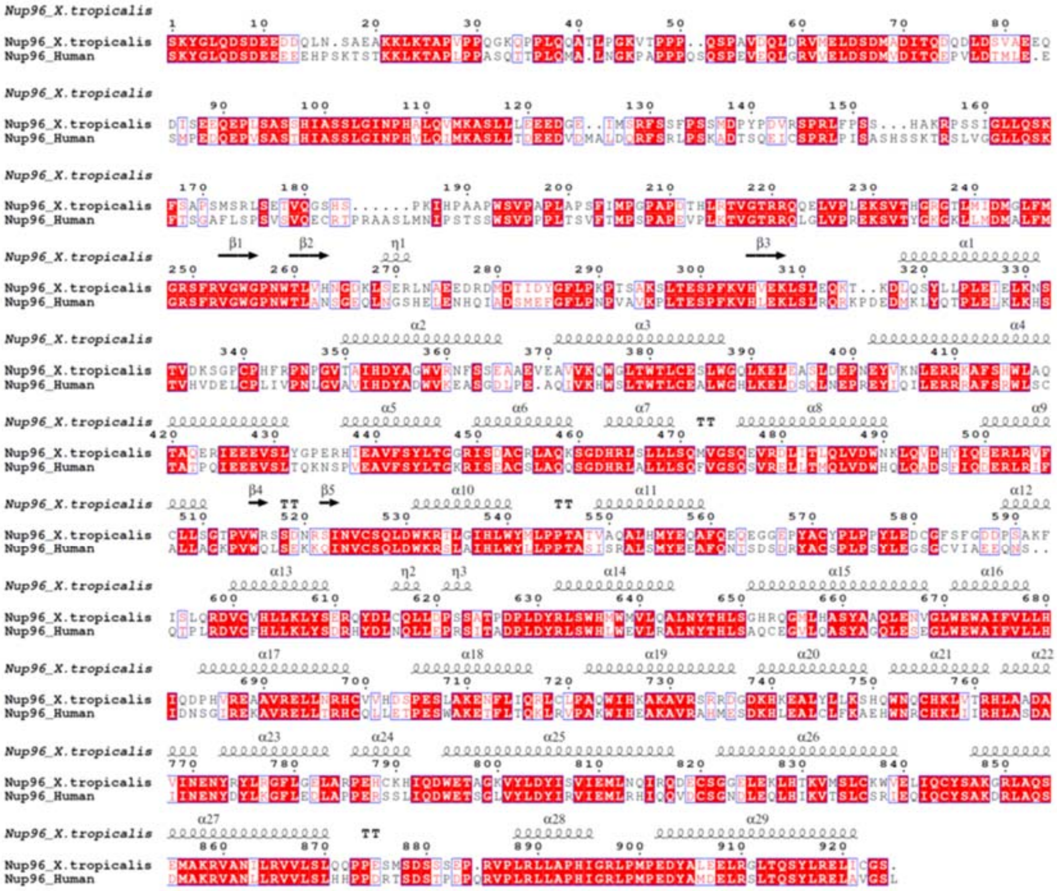

G

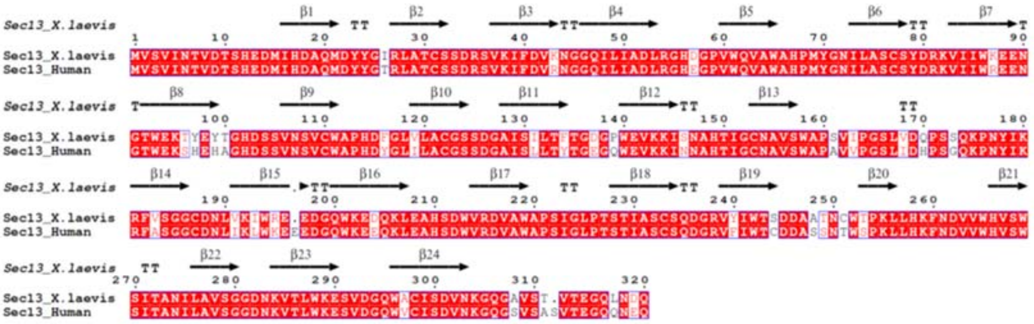


H

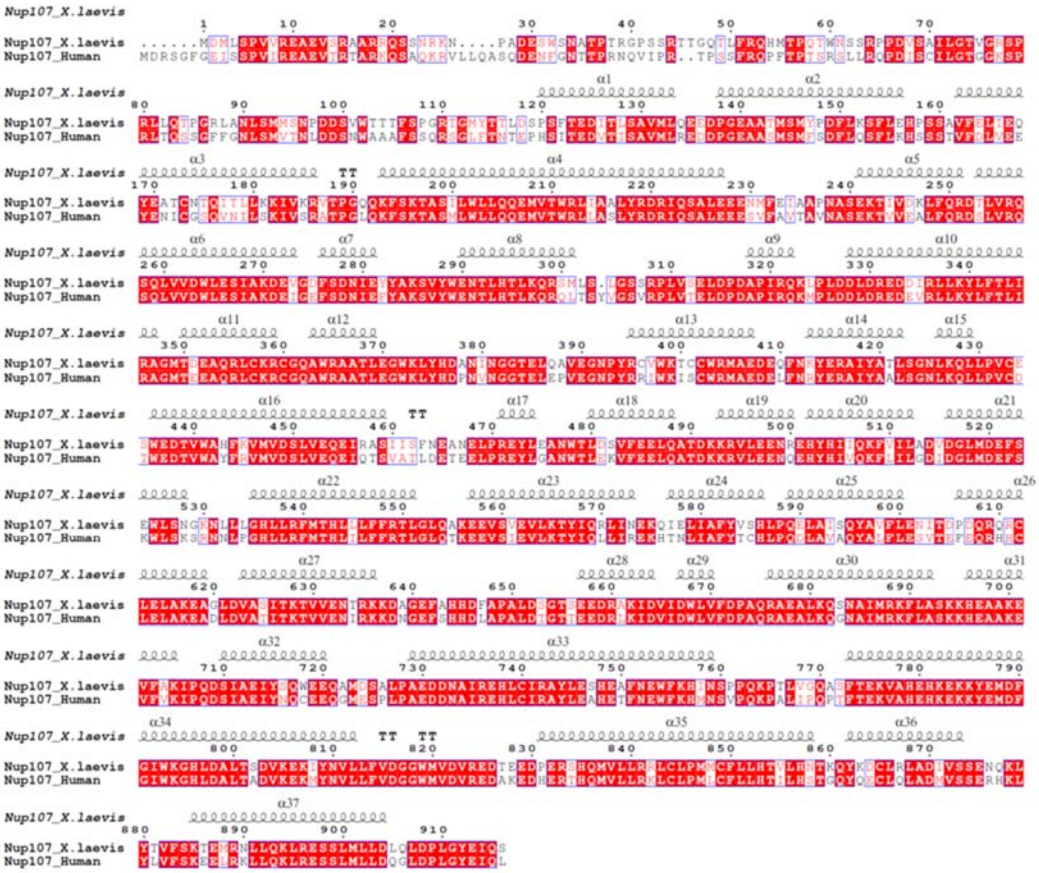

I

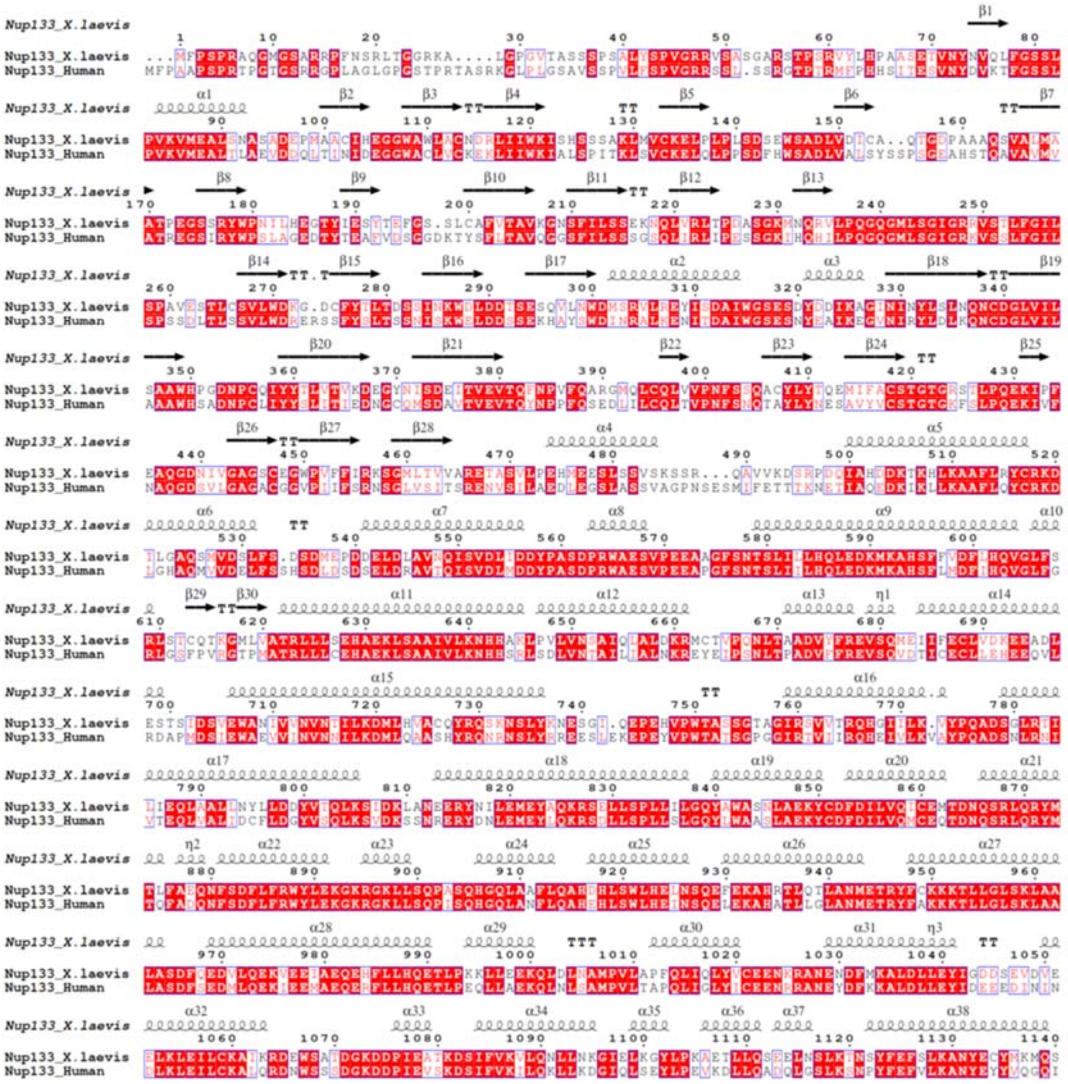

J

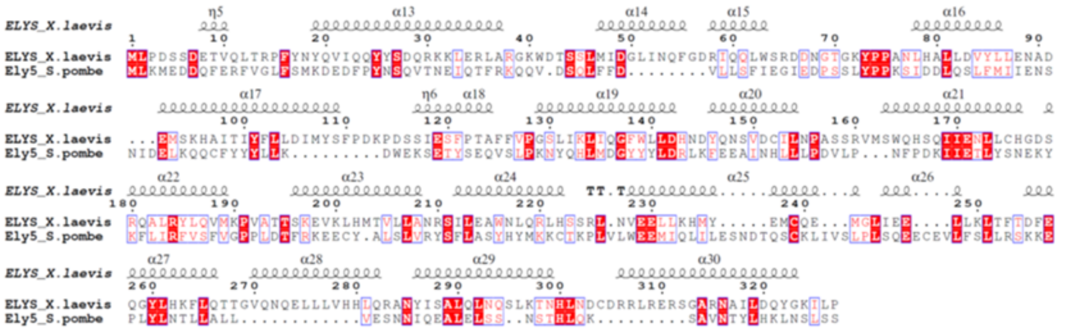

K

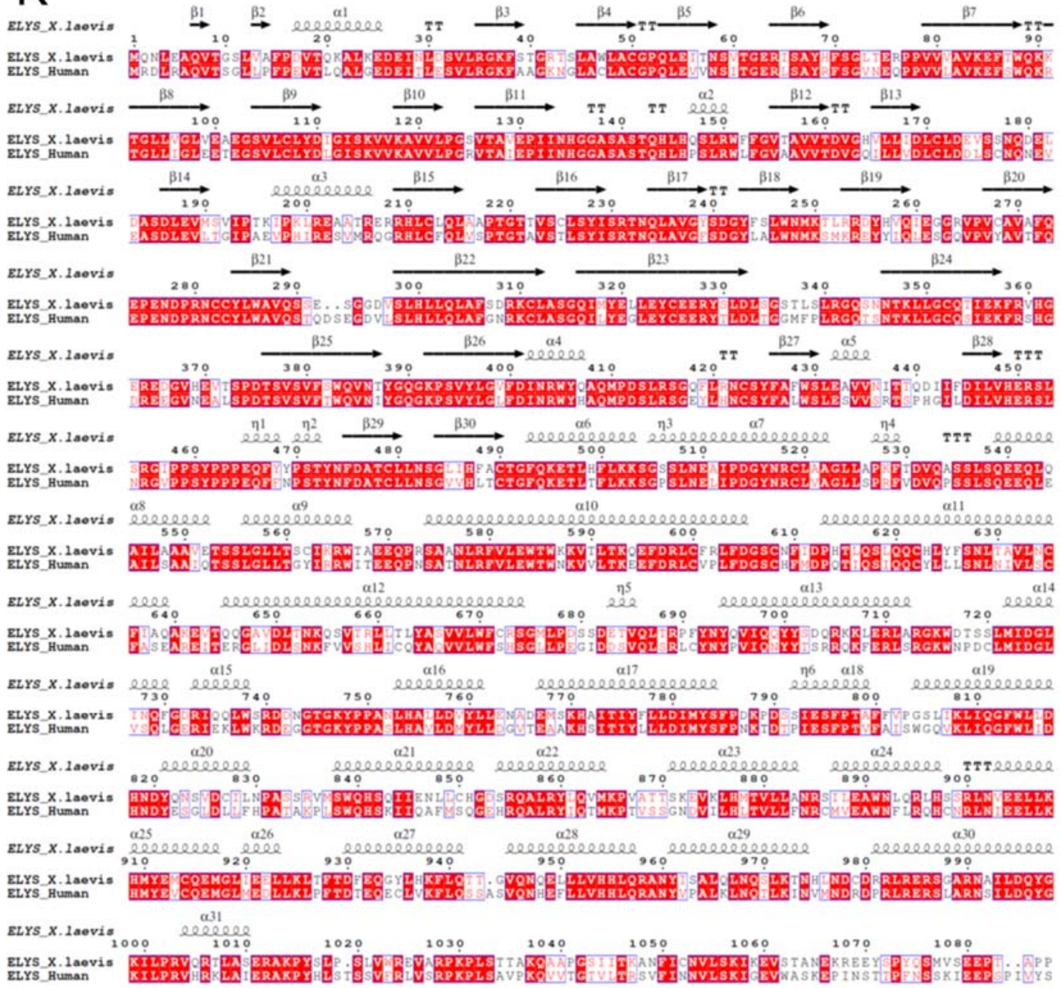


L

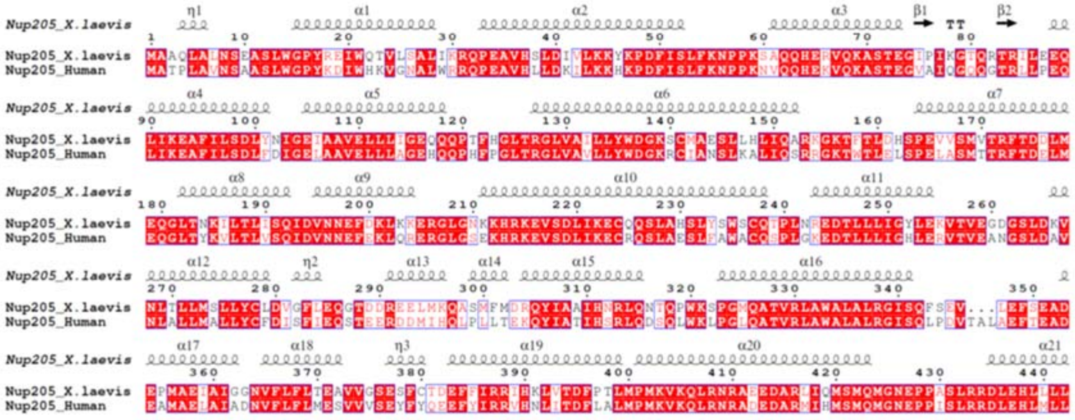

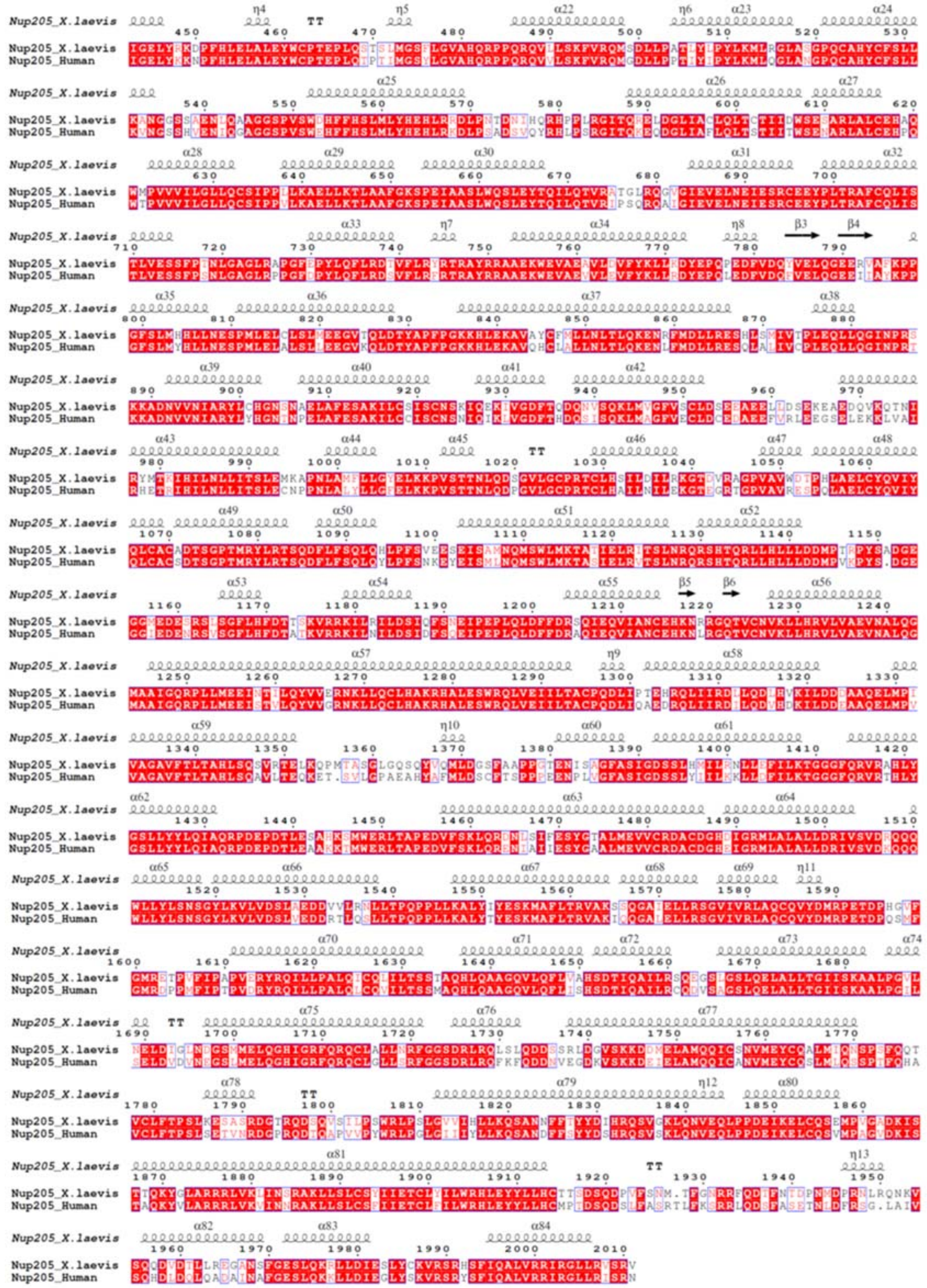

M

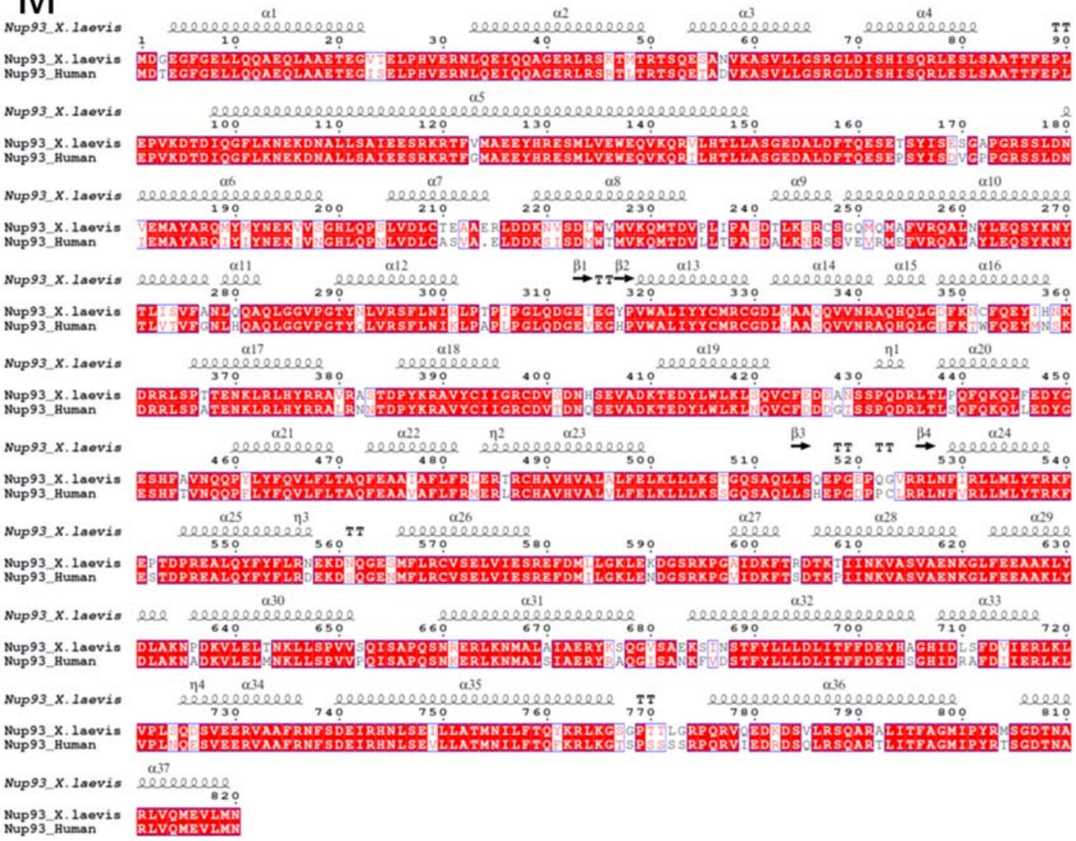


N

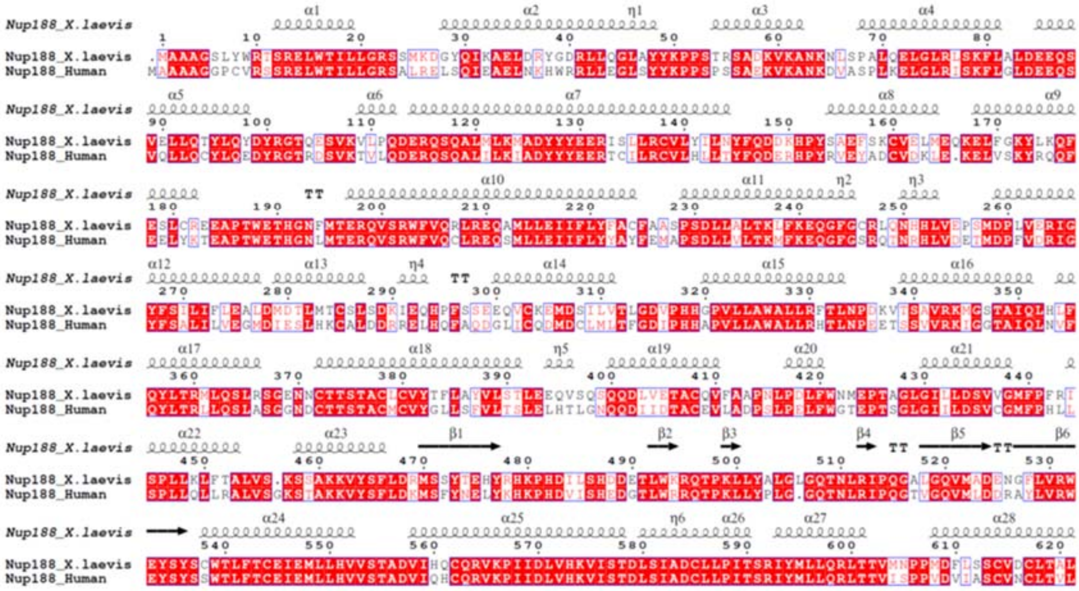

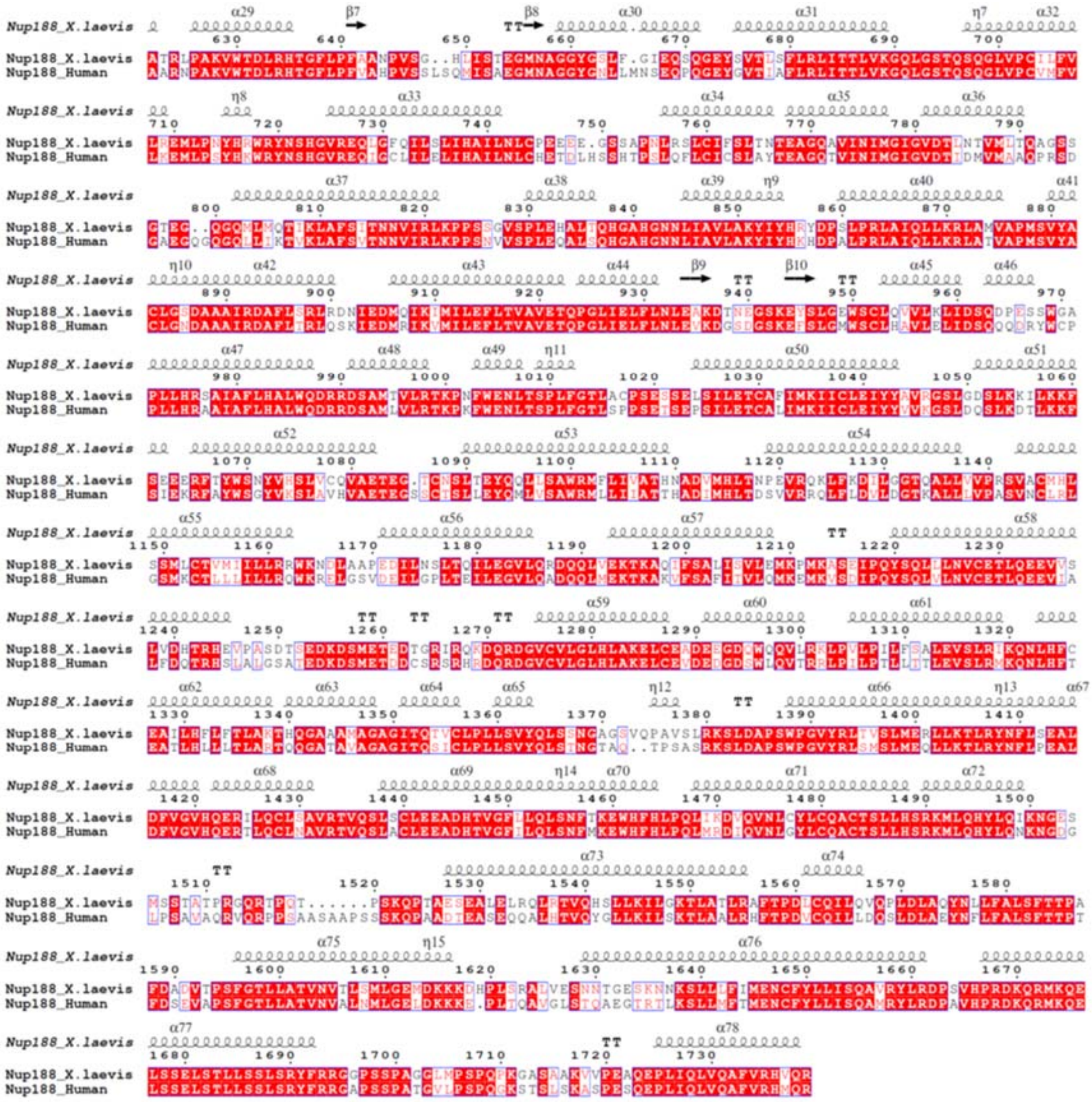

**Extended Data Figure 6. Sequence alignment of NR Nups from *X. laevis* and human NPCs. (A-N)** Nup85, Seh1, Nup43, Nup160, Nup37, Nup96, Sec13, Nup107, Nup133, ELYS, Nup205, Nup93 and Nup188, respectively. Secondary structures are also indicated.

**Extended Data Table 1. Statistics of cryo-SPA data collection and image processing.**

| **Data acquisition** | | | |
|---|---|---|---|
| Microscope | Titan Krios G2 | | |
| Voltage (kV) | 300 | | |
| Detector | Gatan K2 | | |
| Energy filter | Gatan GIF Quantum, 20 eV | | |
| *Mode* | *Super resolution* | | |
| Pixel size (Å) | 2.24 | | |
| Stage tilting angle | 30°/ 45°/ 60°/ 0°(side-view) | | |
| Exposure per tilt (e/Å²) | 60 / 80/ 100/ 100 (or 120) | | |
| *Number of images* | *8745* | | |
| Defocus range (μm) | -1 ~ -4 | | |
| Software | SerialEM | | |
| **Reconstruction** | | | |
| Software | RELION-3.0/CryoSPARC | | |
| Data set | NR core region | NR subunit region | NR Nup133 region |
| Final number of particles | 417490 | 417490 | 298318 |
| Symmetry | C1 | C1 | C1 |
| Final resolution (Å) | 7.8 | 8.1 | 8.6 |
| Map pixel size (Å) | 2.24 | 2.24 | 2.24 |
| Map sharpening B-factor (Å²) | -433 | -541 | -612 |

### Supplementary information

**Supplementary Video 1.**

Model-map fitting quality of NR Nups from the *X. laevis* NPC.

**Supplementary Video 2.**

Structural comparison of Nup93 between the refined model and the model predicted by AlphaFold2.
